# Deep multi-omic profiling reveals extensive mitochondrial remodeling driven by glycemia in early diabetic kidney disease

**DOI:** 10.1101/2023.10.26.564228

**Authors:** Cesare Granata, Vicki Thallas-Bonke, Nikeisha J. Caruana, Kevin Huynh, Cheng Xue Qin, Adrienne Laskowski, Matthew Snelson, Jarryd Anthonisz, Edwina Jap, Georg Ramm, Mark E. Cooper, Peter J. Meikle, David A. Stroud, Rebecca H. Ritchie, Melinda T. Coughlan

## Abstract

Changes in mitochondrial energy metabolism are thought to be central to the development of diabetic kidney disease (DKD); however, whether this response is explicitly driven by systemic glucose concentrations remains unknown. Here, we show that titrating blood glucose concentrations *in vivo* directly impacts mitochondrial morphology and bioenergetics and remodels the mitochondrial proteome in the kidney in early DKD. Mitoproteomic analysis revealed profound metabolic disturbances induced by severe hyperglycemia, including upregulation of enzymes involved in the TCA cycle and fatty acid metabolism, enhanced ketogenesis as well as dysregulation of the mitochondrial SLC25 carrier family. Untargeted metabolomics and lipidomics confirmed the enrichment of TCA cycle metabolites, an increase in triglyceride concentrations, and extensive and specific cardiolipin remodeling. Lowering blood glucose to moderate hyperglycemia stabilized all three omic landscapes, partially prevented changes in mitochondrial morphology and bioenergetics, and improved kidney injury. This study demonstrates altered substrate utilization and energy generation in the kidney early in diabetes, during moderate and severe hyperglycemia and provides new insights into kidney metabolism, which has implications for therapeutic strategies aiming at the reinvigoration of mitochondrial function and signaling in diabetes.

## Introduction

Diabetes is a global healthcare problem, affecting more than 400 million people worldwide with prevalence projected to increase dramatically over the next few decades (Sun et al., 2021). Diabetes is a leading risk factor for disease burden, substantially increasing the risk of heart disease, stroke, cognitive decline and kidney disease. Up to 40% of individuals with diabetes develop microvascular complications, including diabetic kidney disease (DKD). DKD is a progressive disease, and whilst the majority of patients die from cardiovascular events, many will progress to end stage renal disease, which requires renal replacement therapy or dialysis. Since current clinical therapies can only postpone, but not prevent renal disease progression, there is a critical need to identify the pathogenic factors that lead to the onset and progression of DKD in order to develop novel therapeutic targets.

The kidney has a high resting metabolic rate (Wang et al., 2010) and is highly dependent on ATP production by oxidative phosphorylation (OXPHOS) to drive the process of proximal tubular reabsorption to maintain renal function (Soltoff, 1986). Accordingly, the kidney has the second highest content of mitochondria and oxygen consumption after the heart (O’Connor, 2006; Pagliarini et al., 2008). Changes in mitochondrial energy metabolism, including defective respiratory chain function are central to the development of DKD (Higgins and Coughlan, 2014). In diabetes, it is thought that the kidneys are unable to compensate for the increased rate of ATP consumption (Nangaku, 2006; Singh et al., 2008). Multiple studies in experimental diabetes have shown that mitochondrial homeostasis is disrupted, with bioenergetic defects and respiratory chain uncoupling (Che et al., 2014; Coughlan et al., 2009; Daehn et al., 2014; Forbes et al., 2008; Hall and Unwin, 2007; Rosca et al., 2005; Sharma et al., 2013; Sivitz and Yorek, 2010), as well as increased mitochondrial fragmentation (Galloway et al., 2012; Wang et al., 2012; Zhan et al., 2015). Emerging animal data show that a defect in mitochondrial respiratory function may be a primary cause of kidney disease, with a decrease in the ATP pool observed in the diabetic setting prior to evidence of renal injury, and increased oxygen consumption later in the progression of DKD, resulting in intra-renal tissue hypoxia (Coughlan et al., 2016b). In adolescents with type 1 diabetes, increased relative renal hypoxia has been shown, stemming from a potential metabolic mismatch between increased renal energy expenditure due to increased glomerular filtration rate and impaired substrate utilization (Vinovskis et al., 2020).

Whether these mitochondrial adaptations are driven explicitly by hyperglycemia has not been established. To investigate the direct impact of blood glucose on mitochondrial flexibility, we used a well-established model of DKD, the streptozotocin-induced diabetic rat, which displays hallmarks of early but progressive human DKD including albuminuria, hyperfiltration and renal structural injury (Coughlan et al., 2016a). We modulated blood glucose levels using insulin therapy, studied the early changes in renal injury and investigated whether blood glucose trajectories were a key determinant of adaptive changes in renal metabolism including mitochondrial flexibility, signal transduction and substrate metabolism, using a data-driven exploratory multiomics workflow.

## Results

### Modelling graded glycemia in vivo to examine blood glucose trajectories

Sprague Dawley rats were administered either streptozotocin (STZ), a pancreatic β cell toxin, to induce insulin deficiency and result in hyperglycemia, or vehicle (citrate buffer) to serve as a control group. Hyperglycemic rats were treated with either 6-7 units of insulin per day to achieve moderate blood hyperglycemia (MHG), or 1-2 units of insulin per day resulting in severe blood hyperglycemia (SHG); all rats were followed for eight weeks (Figure 1A). By design, SHG rats manifested blood glucose levels >30 mmol/l (consistent with severe hyperglycemia), which was 1.5-fold greater than MHG rats manifesting blood glucose levels of ∼20 mmol/l (consistent with moderate-hyperglycemia) and 5-fold greater than rats in the control group, manifesting blood glucose levels of ∼5 mmol/l (consistent with normal glycemia; NG) (Figure 1B). This was accompanied by an increase in glycated hemoglobin (HbA_1c_), revealing a worsening of glucose control in both MHG and SHG rats (Figure 1C). Plasma C-peptide levels, indicative of endogenous insulin production, were decreased in both groups of hyperglycemic rats when compared to NG (Figure 1D). Body weight was decreased in SHG rats (Figure 1E). Kidney enlargement was evident in both groups of diabetic rats reflecting renal hypertrophy; however, this was significantly greater in SHG compared to MHG rats (Figure 1F).

**Figure 1.**
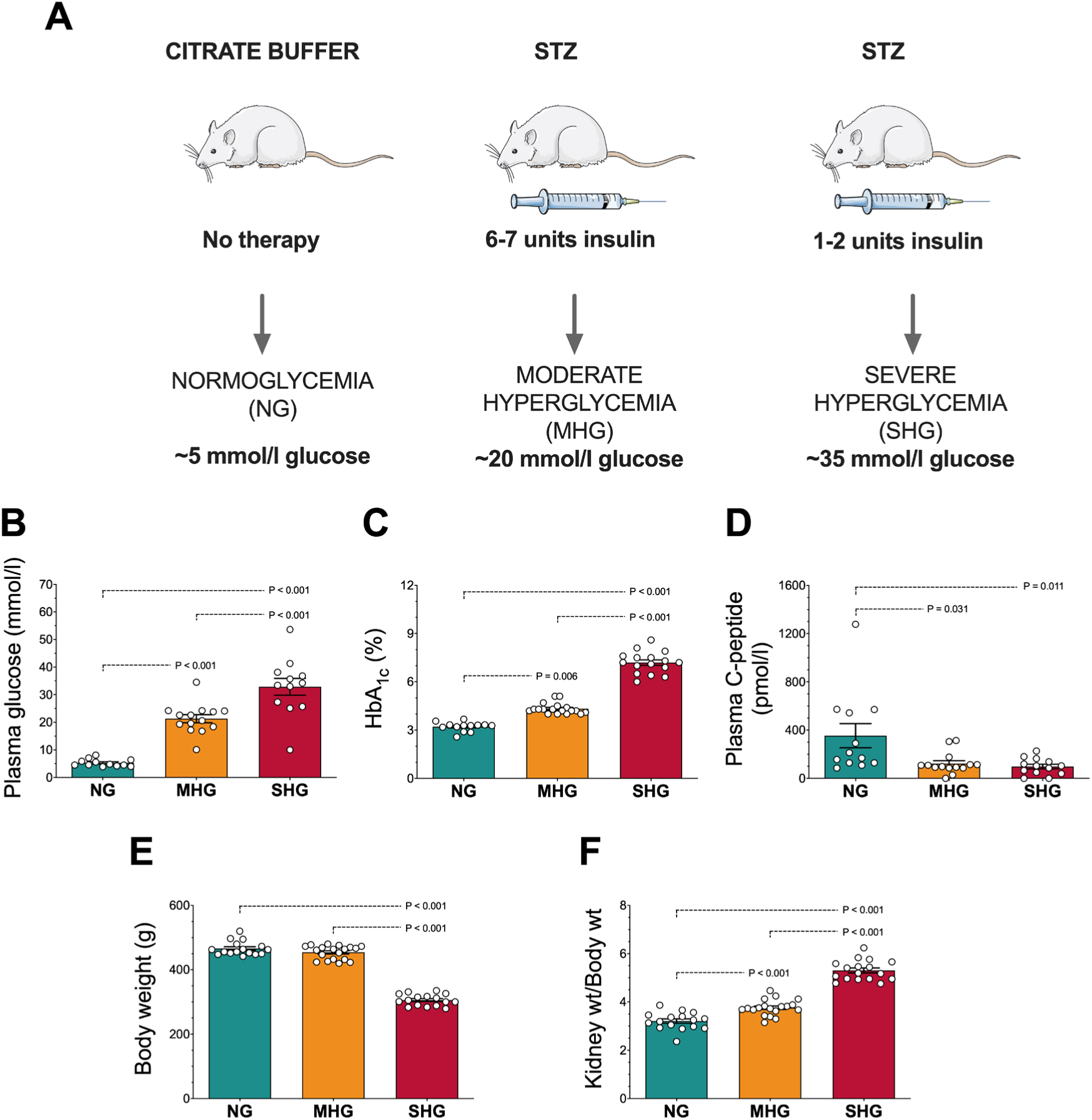
Modelling graded glycemia *in vivo*. (A) Experimental model; (B) Plasma glucose; (C) Hemoglobin A_1c_ (HbA_1c_); (D) plasma C-peptide; (E) body weight; (F) kidney to body weight ratio. Outliers were first removed using the ROUT test with Q = 1%; normality was tested with a Shapiro-Wilk test; data in B and F were analyzed by one way ANOVA followed by Tukey post hoc test; data in C,D, and E were analyzed by Kruskal-Wallis test followed by Dunn’s post hoc test, as not normally distributed; n = 12-14 (B,D), 13-18 (C), 16-19 (E, F). Data are mean ± SEM. Dots represent individual data points. STZ: streptozotocin.

### Reducing hyperglycemia improves the renal phenotype

SHG rats exhibited glomerular hyperfiltration (measured by the surrogate marker of glomerular filtration rate [GFR], cystatin C) (Figure 2A), whereas MHG maintained glomerular filtration at levels similar to NG. Albuminuria was observed in both MHG and SHG rats (Figure 2B). Both levels of hyperglycemia led to morphological changes to the glomeruli of the kidneys, (glomerulosclerosis, Figure 2C and D), indicating that neither insulin dose prevented the hallmarks of DKD, albuminuria and glomerulosclerosis. Tubulointerstitial fibrosis is a prominent feature of progressive chronic kidney disease; tubular injury measured by urinary kidney injury molecule-1 (KIM-1) was evident only in SHG rats (Figure 2E). Similarly, the development of renal fibrosis, as assessed by transforming growth factor-β1 (TGF-β1) (Figure 2F) and collagen IV protein (Figure 2G & H) in the kidney cortex, was only present in SHG rats, indicating that intensive insulin therapy may prevent, or delay, tubulointerstitial fibrosis.

**Figure 2.**
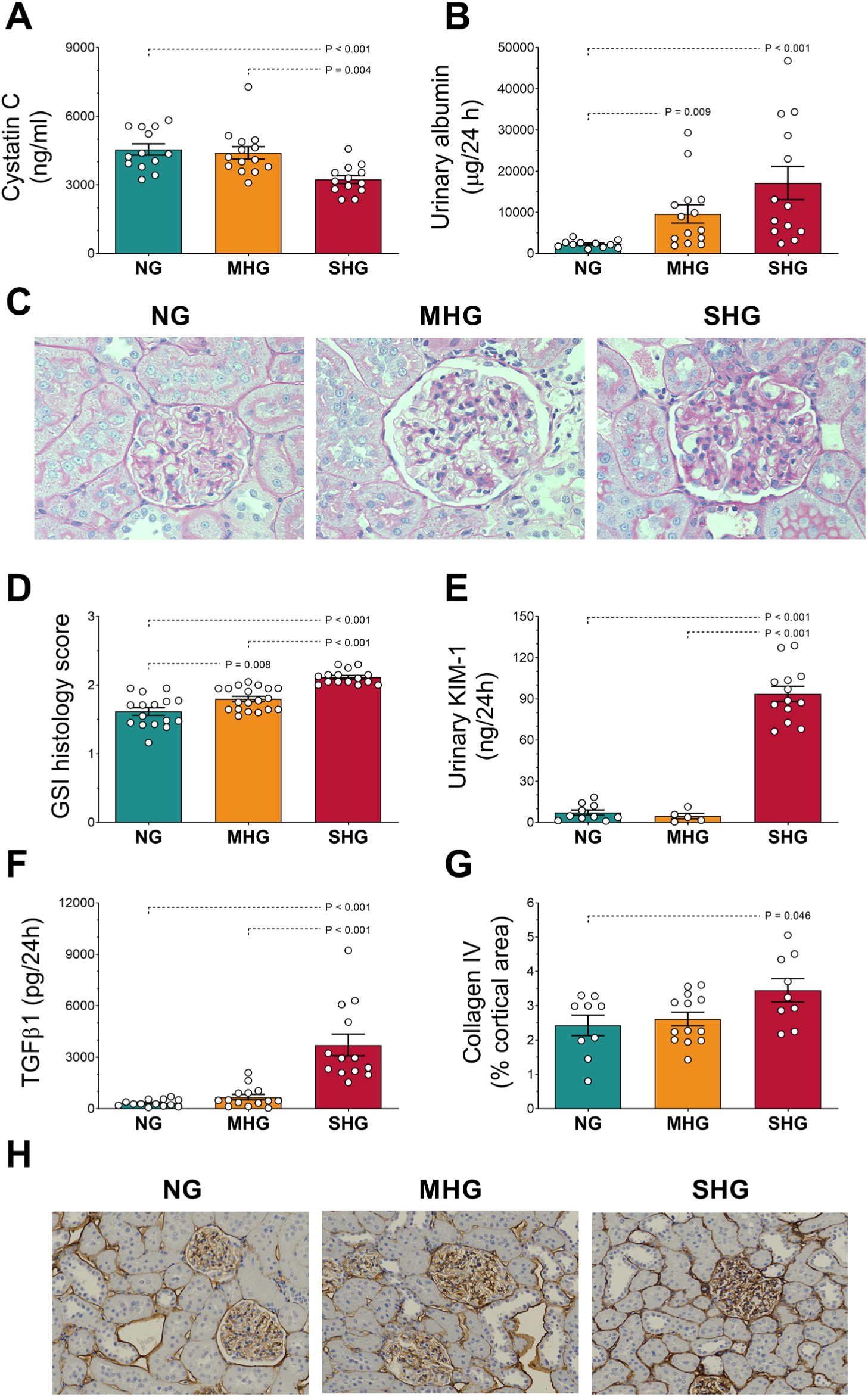
Renal phenotypic changes are improved by lowering blood glucose. (A) Plasma cystatin C; (B) urinary albumin; (C) representative photomicrographs of PAS-stained kidney cortex, x400 magnification related to (D); (D) glomerulosclerotic index (GSI) histology score; (E) urinary excretion of kidney injury molecule-1 (KIM-1); (F) urinary transforming growth factor β (TGFβ1); (G) collagen IV immunohistochemistry of kidney cortex; (H) representative images. NG: normoglycemia; MHG: moderate hyperglycemia; SHG: severe hyperglycemia. Outliers were first removed using the ROUT test with Q = 1%; normality was tested with a Shapiro-Wilk test; data in D, E, and G were analyzed by one way ANOVA followed by Tukey post hoc test; data in A, B, and F were analyzed by Kruskal-Wallis test followed by Dunn’s post hoc test, as not normally distributed; n = 12-14 (A, F), 13-14 (B), 16-19 (D), 6-13 (E), 9-13 (G). Data are mean ± SEM. Dots represent individual data points.

### Hyperglycemia drives defective oxidative phosphorylation, altered mitochondrial morphology, and increased reactive oxygen species (ROS) generation in proximal tubular mitochondria

We next explored mitochondrial respiratory function in the kidney cortex, which is the site of kidney injury in DKD. Citrate synthase (CS) enzyme activity, a marker of mitochondrial content (Larsen et al., 2012), was increased by both grades of hyperglycemia (Figure 3A). Accounting for the increase in mitochondrial content observed by CS, both MHG and SHG led to a significant decrease in the enzymatic activity of OXPHOS complex I (CI, Figure 3B). Measurement of H_2_O_2_ by the Amplex red probe indicated that SHG increased glutamate-malate (GM)-linked ROS generation compared to NG and MHG (Figure 3C), and that both levels of hyperglycemia induced greater succinate-rotenone (SR)-linked ROS generation compared to normoglycemia (Figure 3D). We next assessed mitochondrial morphology by transmission electron microscopy (TEM) within proximal tubule epithelial cells (PTECs). Not only is this cell type dominant within the kidney cortex in terms of cell abundance (Balzer et al., 2022), in human DKD, it displays a substantial injury-associated expression signature (Wilson et al., 2022). TEM images revealed that SHG induced an increase in mitochondrial volume density (Mito_VD_; Figure 3E), as well as a decrease in form factor (an index of mitochondrial complexity and branching (Koopman et al., 2006); Figure 3F) and mitochondrial aspect ratio (length-to-width ratio; Figure 3G), indicating more fragmented mitochondria with less mitochondrial branching. This was supported by a decrease in MFN2 observed in our proteomic analysis (discussed in detail below), an essential protein involved in mitochondrial fusion (Detmer and Chan, 2007). MHG also led to a less branched mitochondrial phenotype as shown by a decrease in form factor (Figure 3F) and an increase in Mito_VD_ that was just short of significance (Figure 3E). Collectively, this indicates that severe hyperglycemia is associated with increased mitochondrial fragmentation and an overall less complex mitochondrial structure, as also indicated by increased circularity and roundness (Figure S1A and B, respectively), whereas moderate hyperglycemia did not display the same severity in these morphological changes.

**Figure 3.**
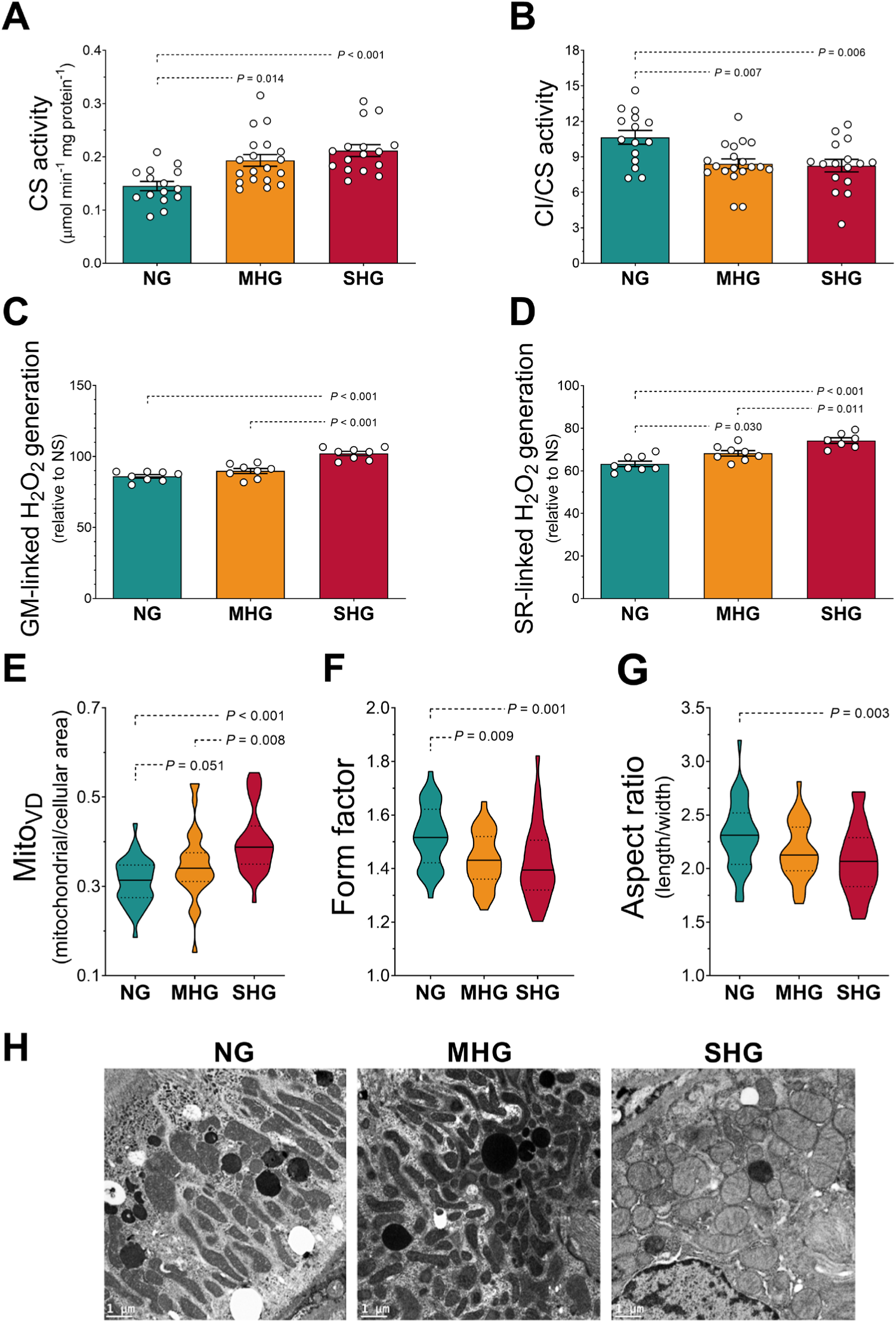
OXPHOS enzyme activity, ROS generation and mitochondrial morphology are perturbed in early DKD and are in part glucose-dependent. (A) Citrate synthase (CS) enzyme activity; (B) complex I (CI) enzyme activity normalized by CS, (C) mitochondrial glutamate-malate (GM)- and (D) succinate-rotenone (SR)-linked H_2_O_2_ generation relative to no-substrate (NS) control; n = 15-19 (A, B), 8 (C, D). Data are mean ± SEM. Dots represent individual data points. (E) Mitochondrial volume density (Mito_VD_); (F) mitochondrial form factor; (G) mitochondrial aspect ratio; (H) representative electron micrographs of renal proximal tubule epithelial cells (PTECs); x15,000 magnification; scale bar = 1 µm; n = 3-4 rats/group and 5-15 images/rat (E, F, G). Bold solid lines represent the median; lighter dashed lines represent the 25 and 75 percentile (interquartile range). NG: normoglycemia; MHG: moderate hyperglycemia; SHG: severe hyperglycemia. Outliers were first removed using the ROUT test with Q = 1%; normality was tested with a Shapiro-Wilk test; data in B, C, D, F, and G were analyzed by one way ANOVA followed by Tukey post hoc test; data in A and E were analyzed by Kruskal-Wallis test followed by Dunn’s post hoc test, as not normally distributed.

### Aberrant glucose control drives changes to the kidney mitochondrial proteome

We next sought to map the changes in the mitochondrial proteome reflective of the blood glucose trajectories. Proteomic analysis was performed on mitochondria isolated from the kidney cortex, resulting in the identification of 2620 proteins (Table S1), 569 of which were annotated as mitochondrial proteins based on the Integrated Mitochondrial Protein Index (IMPI; Known Mitochondrial, reflecting experimental evidence for mitochondrial localization) database (Smith and Robinson, 2019) (Table S2). Multidimensional scaling analysis of mitochondrial proteins readily segregated SHG samples from MHG and NG (Figure 4A). This indicates that SHG induced a distinct differential landscape in the mitochondrial proteome compared to MHG and NG, suggesting that a more severe blood glucose trajectory is associated with extensive mitoproteome remodeling.

**Figure 4.**
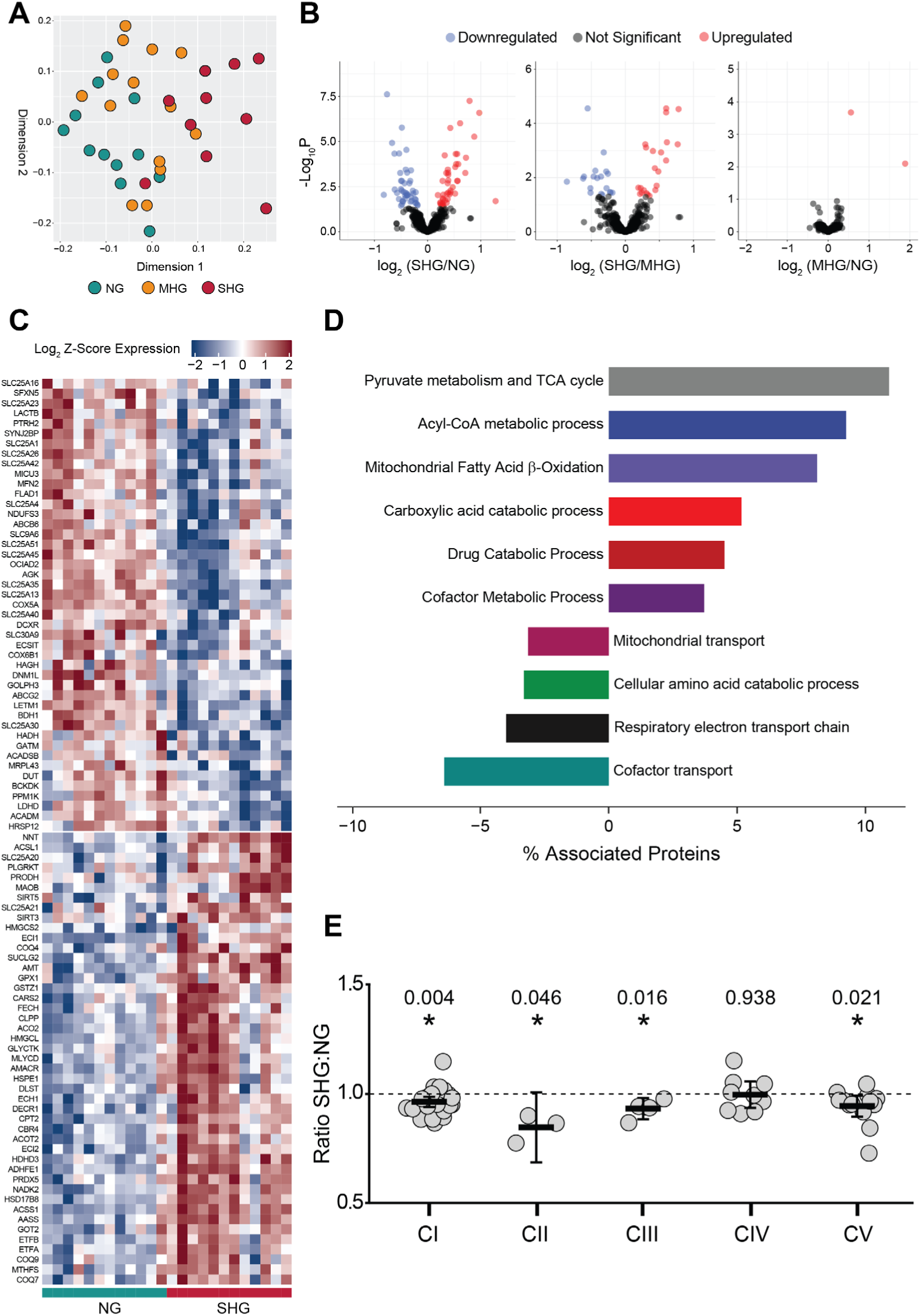
Titrating glucose concentrations remodels the kidney mitoproteome landscape. (A) Multidimensional scaling analysis showing the three treatment groups. (B) Volcano plot representations (unpaired *t*-test) showing differentially expressed proteins between group pairs following permutation-based false discovery rate (FDR) correction; black dots, not significant (*P* > 0.05); blue dots, significant decrease (*P* < 0.05); red dots, significant increase (*P* < 0.05). (C) Unsupervised hierarchical clustering heat map analysis of the 90 differentially expressed mitochondrial proteins between SHG and NG. (D) GO term analysis using ClueGO with GOFusion showing percentage of associated genes for each term for both the positive and negative differentially expressed proteins in SHG *vs.* NG. (E) Analysis of OXPHOS complex protein levels assessed by quantitative proteomics in kidney cortex. Each point represents ratio of the mean of SHG and NG samples for each subunit of each OXPHOS complex, demonstrating the overall change for that complex, n=3-12/group. A * indicates a complex that was significantly reduced in SHG compared to NG (*P* < 0.05), as identified by a Student’s *t*-test; middle bars represents the mean value, while the upper and lower bars represent the 95% confidence interval of the mean value. Each dot represents a single subunit. All other panels in the figure: n = 12/group. NG: normoglycemia; MHG: moderate hyperglycemia; SHG: severe hyperglycemia.

Further analyses revealed that 76 mitochondrial proteins were differentially expressed across all three treatment groups (Table S3). To better characterize differences in the mitoproteome, we performed direct group comparisons and identified 90, 41, and two differentially regulated mitochondrial proteins when comparing the SHG *vs.* NG, SHG *vs.* MHG, and MHG *vs.* NG groups, respectively (Figure 4B; Table S3). Of the 41 differentially expressed proteins identified between SHG and MHG, 37 were in common with the SHG *vs.* NG comparison (Figure S2); the direction of change (up- or down-regulation) was also maintained. This suggests that proteomic changes between the two hyperglycemic groups were similar, albeit less pronounced, to changes occurring between SHG and NG. Conversely, only two mitochondrial proteins, HMGCS2, a key enzyme involved in the biosynthesis of ketone bodies (Hegardt, 1999), and ECI1, an enzyme involved in the β-oxidation of unsaturated fatty acids (Kunau et al., 1995), were differentially expressed (both upregulated) in MHG *vs.* NG rats (Figure 4B; Table S3), consistent with previous research (Sas et al., 2016; Shukla et al., 2017; Zhang et al., 2010). The minimal changes observed in the kidney mitochondrial proteome of MHG rats compared to NG mirror the prevention of hyperfiltration, tubular injury, and renal fibrosis afforded by the intensive insulin therapy described above. Taken together these results indicate that intensive insulin therapy, resulting in moderate hyperglycemia, was able to prevent, or delay, most diabetes-induced proteomic changes. As a result, our investigation will focus on the comparison between SHG *vs.* NG.

To characterize the changes to the mitochondrial proteome induced by severe hyperglycemia we performed unsupervised hierarchical clustering on the 90 differentially expressed proteins between SHG and NG (Figure 4C). Enrichment analysis revealed that SHG induced changes in proteins participating in many central mitochondrial metabolic pathways compared to NG (Figure 4D; Table S4). These included an increase in proteins involved in, amongst others, pyruvate metabolism and the tricarboxylic acid (TCA) cycle, as well as acetyl CoA metabolic process and mitochondrial fatty acid β-oxidation (Figure 4D; Table S4). Conversely, there was a reduction in proteins associated with cofactor transport, mitochondrial transport, and mitochondrial protein import, which was driven, in large part, by extensive reduction in the abundance of proteins belonging to the mitochondrial carrier SLC25 family, which are responsible for transport of small molecules across the inner mitochondrial membrane (Figure 4D; Table S4). Proteins associated with the respiratory electron transport chain (ETC) Gene Ontology (GO) term were also less abundant in SHG, an effect driven by a reduction in respiratory chain complex subunits (discussed later). Taken together, these findings indicate that severe hyperglycemia induces extensive remodeling of the mitochondrial proteome, with strong emphasis on nutrient transport and utilization, raising the possibility that these changes may be associated with development of renal injury and mitochondrial dysfunction.

### Severe hyperglycemia induces a profound dysregulation of the mitochondrial carrier family SLC25

We next investigated some of the over-represented pathways highlighted by our enrichment analysis beginning with the pathways related to mitochondrial protein transport (Figure 4D; Table S4). The SLC25 family of solute carriers, which are embedded in the inner mitochondrial membrane (IMM), is involved in the transport of a variety of solutes essential for the function of multiple metabolic pathways (Palmieri, 2013). Our proteomics analysis indicated that the relative abundance of 14 of the 25 identified SLC25s was altered by severe hyperglycemia, with 12 SLC25s being significantly reduced in SHG (Figure 4C; Table S3). Of these, we report a strong SHG-induced decrease in the abundance of SLC25A30 (also known as KMCP1 or UCP6). SLC25A30, which facilitates the export of hydrogen sulfide degradation products outside the mitochondria (Gorgoglione et al., 2019), is a kidney-specific protein involved in the regeneration from tubular injury and in the protection from oxidative damage following increased mitochondrial metabolism (Haguenauer et al., 2005). The decrease in SLC25A30 in SHG is consistent with a reduced ability to protect against the increase in oxidative stress (Figure 3C and D) and a compromised capacity for regeneration of tubular injury (Figure 2E).

### Severe hyperglycemia impairs the OXPHOS system

Our proteomics analysis showed that severe hyperglycemia induced a significant decrease in COX5A and COX6B1, as well as NDUFS3, subunits of respiratory chain CIV and CI, respectively (Figure 4C; Table S3). We calculated changes in the abundance of the OXPHOS complexes using a method we previously established for assessing the impact of OXPHOS defects in patients with suspected mitochondrial disease (Helman et al., 2021; Lake et al., 2017); our results show a significant decrease in the abundance of four of the five OXPHOS complexes (Figure 4E). We also found that LDHD, which catalyzes the oxidation of D-lactate to pyruvate while feeding electrons to the ubiquinone/cytochrome enzyme in the ETC (Kohn and Kaback, 1973), was severely decreased by SHG (Figure 4C; Table S3). These findings, together with the decrease in CI activity (Figure 3B), suggest that OXPHOS is impaired in SHG, consistent with previous reports (Zhang et al., 2018).

As introduced above, our proteomics analysis shows dysregulation of solute carriers, some of which are directly linked to OXPHOS through their roles shuttling adenine nucleotide species across the IMM. We observed a decrease in SLC25A4 (also known as ANT1) (Figure 4C; Table S3), the key solute carrier responsible for ADP/ATP shuttling across the IMM (Klingenberg, 2008). Knockout of this carrier in mice has been linked to decreased CI content and activity (McManus et al., 2019). SLC25A23 (also known as APC2), was also decreased (Figure 4C; Table S3); SLC25A23, together with other APCs, modulates matrix adenine nucleotide-dependent anabolic pathways (e.g., gluconeogenesis, urea synthesis, nuclear-encoded mitochondrial protein import, mtDNA-encoded protein synthesis) and adenine nucleotide concentrations (Palmieri, 2013). SLC25A13, an aspartate/glutamate carrier (also known as AGC2) that is also involved in the urea cycle and in gluconeogenesis (Ruprecht and Kunji, 2019), was also reduced in SHG (Figure 4C; Table S3). Taken together, the above findings provide evidence that severe hyperglycemia results in both dysfunctional OXPHOS and a reduction in anabolic processes.

### Severe hyperglycemia drives an increase in the TCA cycle and a shift away from OXPHOS dependent metabolism

Enrichment analysis indicated a severe hyperglycemia-induced increase in abundance of TCA cycle enzymes (Figure 4D; Table S4), which was underpinned by a significant increase in the protein content of ACO2, DLST and SUCLG2 (Figure 4C; Table S3) (Figure 3A). To further investigate changes in this metabolic pathway, we performed a mass-spectrometry based metabolomics assessment of intracellular metabolites in whole-kidney cortex. Our metabolome analysis identified 165 metabolites (Table S5), 77 of which were differentially expressed between different levels of glycemia (Table S6). Principal component analysis of all identified metabolites segregated, once again, SHG from MHG and NG samples (Figure S3A). Owing to this and to findings from our proteomic analysis indicating that the main differences took place between SHG and NG, we focused our analysis on this comparison and the 70 differentially expressed metabolites identified between SHG and NG (Table S6). Unsupervised hierarchical clustering of all differentially expressed metabolites between SHG *vs.* NG showed clear segregation of samples with different levels of glycemia in two main clusters with opposite features (Figure 5A; Table S6). Enrichment analysis indicated SHG-induced increases in metabolites involved in the central carbon metabolism, TCA cycle, the Warburg effect, the transfer of acetyl groups into mitochondria, and glycolysis and a decrease in metabolites involved in homocysteine degradation, catecholamine biosynthesis, biotin metabolism, glycine and serine metabolism, and methionine metabolism (Figure 5B; Table S7). As a consequence of the above findings, we performed an in-depth analysis of all the identified metabolites of the TCA cycle; this showed a large (approximately two-fold) and concerted increase in all metabolites involved, with the exception of oxaloacetate, which remained unchanged (Figure 5C).

**Figure 5.**
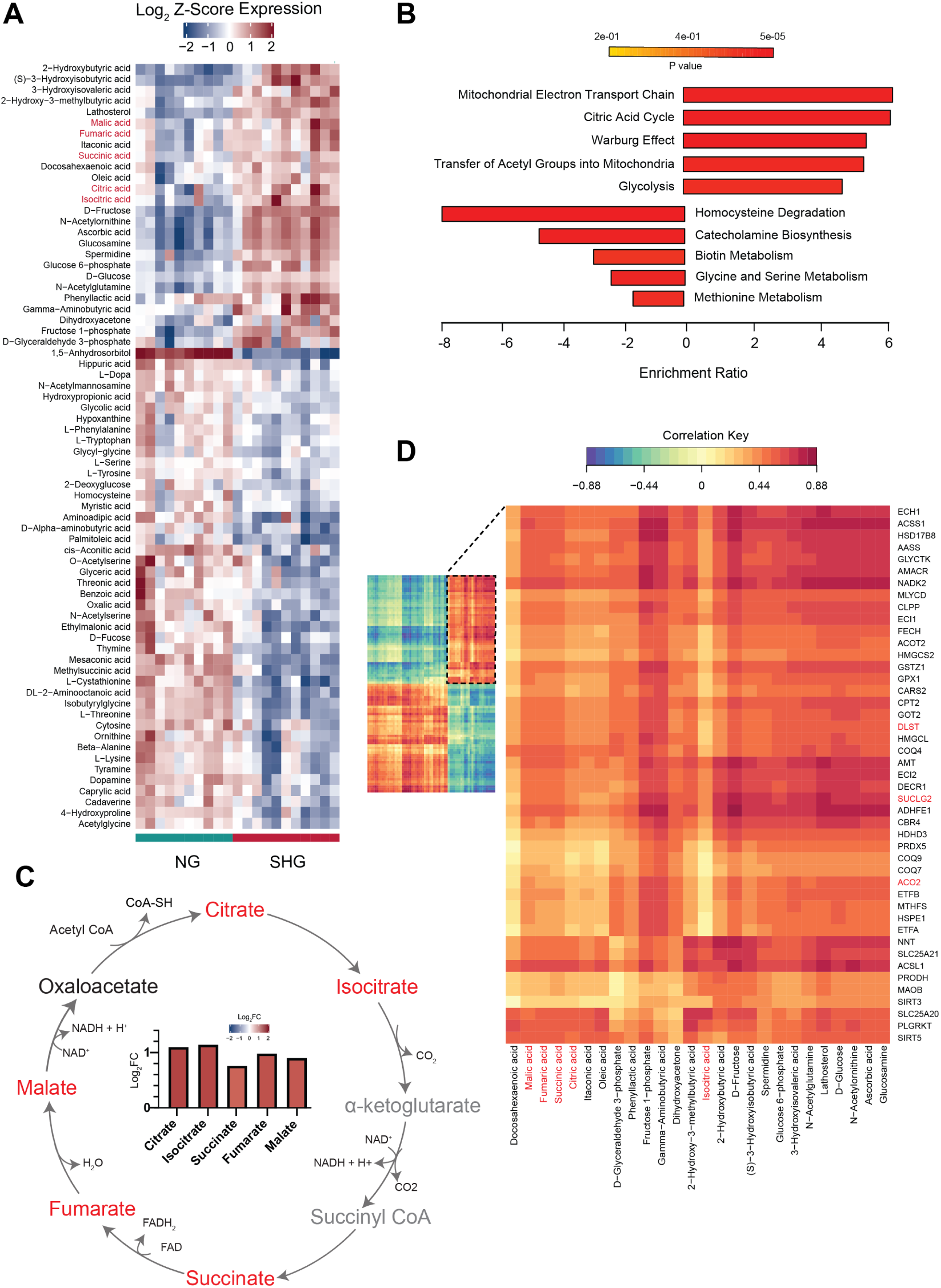
Metabolomics reveals distinct changes in substrate utilization and energy generation. (A) Unsupervised hierarchical clustering heat map analysis of the 70 differentially expressed metabolites between SHG and NG; TCA cycle metabolites are highlighted in red. (B) Overrepresentation analysis showing the enrichment ratio for the top 5 overrepresented upregulated and downregulated pathways from the SHG *vs.* NG comparison, run independently in Metaboanalyst. (C) Tricarboxylic acid (TCA) cycle representation showing unchanged (black), increased (red), and not identified (grey) metabolites, with bar graph showing log_2_FC for the significantly increased TCA metabolites (adj. *P* < 0.05). (D) Regularized canonical correlation analysis between the differentially expressed proteins (rows) and metabolites (columns) between SHG vs. NG; TCA cycle metabolites and enzymes are highlighted in red. All panels in the figure: n = 10 (NG) and 11 (SHG). NG: normoglycemia; SHG: severe hyperglycemia.

To further investigate these findings, we performed a regularized canonical correlation analysis between all the differentially expressed proteins (rows) and metabolites (columns) from the SHG *vs.* NG comparison (Figure 5D). As shown in the inset, a strong correlation was observed between the subset of upregulated proteins, which were enriched in TCA cycle enzymes (Figure 4D; Table S4) with the subset of upregulated metabolites, which was also enriched in TCA cycle intermediates (Figure 5B; Table S6 and S7; Figure 5D, with upregulated TCA cycle metabolites highlighted in red). These findings suggest that severe hyperglycemia induced an increase in TCA cycle flux. Taken together with the aforementioned impairments in CI activity, this suggests that severe hyperglycemia may promote a metabolic switch towards a more glycolytic phenotype (Warburg effect), a previously suggested feature of DKD (Zhang *et al*., 2018). Opposite to the effects of SHG, only minor changes in abundance of metabolites involved in the TCA cycle and a decrease in cis-aconitate (Table S6) were reported in MHG. This, together with the lack of significant changes reported in proteins involved in the TCA cycle following MHG (Table S3), suggest that intensive insulin therapy (leading to MHG) helps to prevent, or delay, most of the metabolic shift induced by severe hyperglycemia.

The increase in abundance of TCA cycle enzymes and metabolites in SHG was also accompanied by extensive dysregulation of several members of the SLC25 family linked to central carbon metabolism (Figure 4C and D). Notably, we report an increase in SLC25A21, responsible for shuttling α-ketoglutarate across the IMM (Palmieri, 2004), and a decrease in two solute carriers responsible for the transport of citrate out of mitochondria: SLC25A1 (Palmieri, 2004) and SFXN5 (Li et al., 2010). This decrease in abundance of citrate carriers may suggest that under SHG citrate is predominantly used in the matrix for catabolic processes (e.g., TCA cycle, fatty acid β-oxidation), rather than in the cytosol for anabolic reactions (e.g., lipogenesis). Because SLC25A1 inhibition has been observed to normalize hyperglycemia in rat liver (Tan et al., 2020a), the reported decrease in SLC25A1 in the present study may serve as a protective mechanism against SHG in the kidney. Finally, there was also a reduction in SLC25A42 and SLC25A16, which is somewhat unexpected, as their main role is to import CoA in the mitochondrial matrix (Fiermonte et al., 2009; Prohl et al., 2001) for use in catabolic processes such as the TCA cycle and β-oxidation, both of which were markedly increased in SHG. Further research is needed to better characterize the remodeling induced by severe hyperglycemia in these solute carriers.

### Severe hyperglycemia induces upregulation of the carnitine shuttle and other enzymes involved in transport of fatty acids to mitochondria for β-oxidation

The carnitine shuttle consists of 3 enzymes required for long-chain fatty acids to cross the inner mitochondrial membrane (IMM) for subsequent β-oxidation. Our results indicate that severe hyperglycemia induced an increase in the abundance of SLC25A20, the solute carrier responsible for the transport of long chain acyl-carnitine substrates across the IMM, and CPT2, the enzyme releasing acyl-CoA from acyl-carnitine in the mitochondrial matrix (Figure 4C; Table S3). CPT1B, one of the two isoforms of the third enzyme involved in this shuttle, also tended to increase, but fell short of statistical significance (Log_2_FC = 0.480, adjusted *P* = 0.071; Table S3). Conversely, CPT1A, whose overexpression has been shown to protect from renal fibrosis by restoring mitochondrial homeostasis (Miguel et al., 2021), remained unchanged (Log_2_FC = 0.012; *P* = 0.886; Table S3). Moreover, our lipidomics analysis in isolated mitochondria from kidney cortex (results presented below) demonstrated that SHG induced increases in abundance of the long-chain acylcarnitines AC(17:0), AC(18:0), and AC(18:1). Taken together, these findings suggest that severe hyperglycemia induced an increase in the carnitine shuttle in rat kidney.

ACSL1, an outer mitochondrial membrane (OMM) enzyme responsible for the activation of long-chain (10 to 20 carbon atoms) fatty acids to acyl-CoA (Waku, 1992), was also increased in SHG (Figure 4C; Table S3). Formation of acyl-CoA by ACSL1 represents the first step of several lipid metabolic processes, both anabolic and catabolic, with knockout studies demonstrating that ACSL1 acts as a metabolic rheostat directing lipids inside the mitochondria for β-oxidation (Young et al., 2018). The increase in ACSL1 in the present study further highlights that severe hyperglycemia induced funneling of lipid species inside the mitochondria for lipid catabolism.

### Severe hyperglycemia upregulates enzymes involved in the β-oxidation of unsaturated fatty acids, and decreases branched-chain amino acid and fatty acid catabolism

Under physiological conditions, PTECs depend largely on lipolysis and fatty acid β-oxidation for ATP production (Forbes and Thorburn, 2018), with previous literature demonstrating that dysregulation of lipid metabolism is an important factor associated with DKD (Stadler et al., 2015). In line with this, and consistent with the upregulation of the carnitine shuttle, our proteomic enrichment analysis indicated an upregulation of lipid metabolism in SHG (Figure 4D; Table S4). Specifically, we identified an increase in several auxiliary enzymes involved in fatty acid β-oxidation; these included ECI1 and ECI2, ECH1, and DECR1 (Figure 4C; Table S3), which are enzymes involved in the β-oxidation of mono-, di-, and poly-unsaturated fatty acids (Schulz and Kunau, 1987). These findings suggest that severe hyperglycemia resulted in an increase in lipid catabolism in kidney cortex that was likely driven by upregulation of β-oxidation of unsaturated fatty acids.

Mitoproteomics analysis also revealed that the protein content of the medium-chain specific ACADM (ACAD1) and the short and branched-chain (BC) specific ACADSB (ACAD7), two of the ACADs involved in the catalysis of the first step of β-oxidation, were reduced (Figure 4C; Table S3). Whereas the enzymatic activity of ACADM can be performed by other ACADs due to the overlap in their affinity for different carbon chain lengths (He et al., 2011), the catalytic activity of ACADSB is specific to BC amino acids (BCAAs) and short (C4 to C6) BC fatty acids (BCFAs) (Rozen et al., 1994). This was accompanied by a concomitant decrease in the content of BCKDK, the kinase responsible for the regulation of the BCKD complex (Harris et al., 1997), which controls BCAA breakdown in several tissues (Holeček, 2018). These findings suggest a decrease in BCAA and short BCFA catabolism, as also indicated by our enrichment analysis (Figure 4D; Table S4), and are consistent with previous reports showing dysregulated BCKD activity in diabetes and renal failure (Holeček, 2018).

The abundance of ETFA and ETFB was also significantly increased (Figure 4C; Table S3). ETFA and ETFB are key recipients of the electron transfer occurring during oxidation of acyl CoAs, and are therefore the primary entry point in the OXPHOS process for the reducing equivalents generated by fatty acid β-oxidation (Pesta and Gnaiger, 2012). The increase in the protein content of these ETFs, coupled with the increase in TCA cycle, indicates that oxidative phosphorylation, although overall downregulated in SHG, may be mainly fueled by reducing equivalents generated during lipolysis. In conclusion, the above findings, together with the upregulation of the carnitine shuttle, provide strong evidence that severe hyperglycemia induces upregulation of lipid catabolic processes in the kidney.

As a corollary to the increase in fatty acid metabolism, the protein content of ACOT2, a protein catalyzing the hydrolysis of acyl-CoAs to the free fatty acid and CoA (Moffat et al., 2014) was also increased by SHG (Figure 4C; Table S3). ACOT2 is a key modulator of intracellular levels of acyl-CoAs, free fatty acids and CoA, and a regulator of subcellular trafficking of fatty acid between mitochondria and peroxisomes (Tillander et al., 2017), which has been implicated in the pathophysiology of type 1 diabetes (King et al., 2007; Moffat *et al*., 2014; Tillander *et al*., 2017). Because lipid accumulation in the kidney is thought to lead to lipotoxicity and kidney injury, we hypothesize that the increase in ACOT2 may orchestrate a protective mechanism by increasing fatty acid and acyl-CoA trafficking between the mitochondria and the peroxisomes, in an attempt to circumvent the toxic effects induced by renal lipid build up, as has been observed in the liver (Moffat *et al*., 2014).

### Severe hyperglycemia is associated with increased ketogenesis

In the kidney, ketone bodies serve as alternative fuels when challenged or during fasting conditions (Forbes and Thorburn, 2018). Owing to this and the increase in fatty acid β-oxidation associated with SHG, we investigated pathways of ketone body metabolism. Our proteomics data show that SHG induced an increase in HMGCS2 and HMGCL (Figure 4C; Table S3), two key ketogenic enzymes generating acetoacetate from acetoacetyl CoA and acetyl CoA (Newman and Verdin, 2014). ACSS1, the enzyme responsible for the activation of acetate to acetyl-CoA, a precursor metabolite central to several metabolic processes including ketogenesis (Pietrocola et al., 2015), was also markedly increase in SHG (Figure 4C; Table S3). This suggests that ketogenesis was increased by severe hyperglycemia, consistent with findings in the kidney of an STZ rat model (Zhao et al., 2011). Meanwhile, acetoacetate levels remained constant in SHG kidneys (Table S6) despite the increase in HMGCS2 and HMGCL, as well as the marked decrease in BDH1 (Table S3), the enzyme catalyzing the interconversion between acetoacetate and β-hydroxybutyrate (Newman and Verdin, 2014). The lack of change in acetoacetate levels may be explained by an increase in cholesterol synthesis in the kidney (Carreau et al., 1982), as suggested by the marked increase in lathosterol (Table S6), a metabolite in the cholesterol synthetic pathway, and/or by its non-enzymatic degradation to acetone (not detectable by our metabolomics method). Moreover, owing to the kidney’s ability to both generate (ketogenesis) and utilize (ketolysis) ketone bodies (Forbes and Thorburn, 2018; Zhang *et al*., 2010), generation of β-hydroxybutyrate (a more stable molecule generated by BDH1 to facilitate delivery to others utilizing tissues (Newman and Verdin, 2014)), might have also been bypassed in an attempt to streamline this metabolic process. OXCT1, a key ketolytic enzyme catalyzing the formation of acetoacetyl-CoA from acetoacetate for subsequent entry into the TCA cycle (Newman and Verdin, 2014), was not changed by SHG (Table S3). Taken together, the above findings indicate that severe hyperglycemia is associated with an increased rate of ketogenesis, but no change in ketolysis in rat kidney cortex.

### Lipidomic analysis identifies hyperglycemia-induced remodeling of mitochondrial cardiolipin

Due to previous findings in mouse tissue indicating a severe diabetes-induced dysregulation of the kidney cortex lipidome (Ge et al., 2020; Sas et al., 2021; Tan et al., 2020b) and the number of enzymes and proteins involved in fatty acid β-oxidation that were differentially expressed by severe hyperglycemia in our study, we next explored the lipidomic landscape in mitochondrial isolates from renal cortex. Our lipidomics analysis identified 666 species (Table S8), 506 of which were differentially expressed between the three levels of glycemia (Table S9). Direct comparison of SHG *vs.* NG, SHG *vs.* MHG, and MHG *vs.* NG resulted in 447, 424, and 137 differentially expressed lipids, respectively (Table S9). Similar to proteomics and metabolomics, multidimensional scaling analysis of all the identified lipids segregated SHG from MHG and NG samples (Figure S3B). Moreover, SHG *vs.* NG changes were more extensive and more pronounced than changes between the other pairs of groups (compare Figure 6A with Figure S4A and B). For these reasons, our lipidomics analysis focused, once again, on the differences between SHG and NG. Unsupervised hierarchical clustering of the 37 identified lipid classes indicated that SHG induced an extensive lipidome dysregulation compared to NG (20 differentially expressed lipid classes; Figure 6B), consistent with previous findings in mouse models of diabetes (Sas *et al*., 2021). The increase in triglycerides (TGs) induced by SHG was one of the predominant features (Figure 6A and B; Table S9).

**Figure 6.**
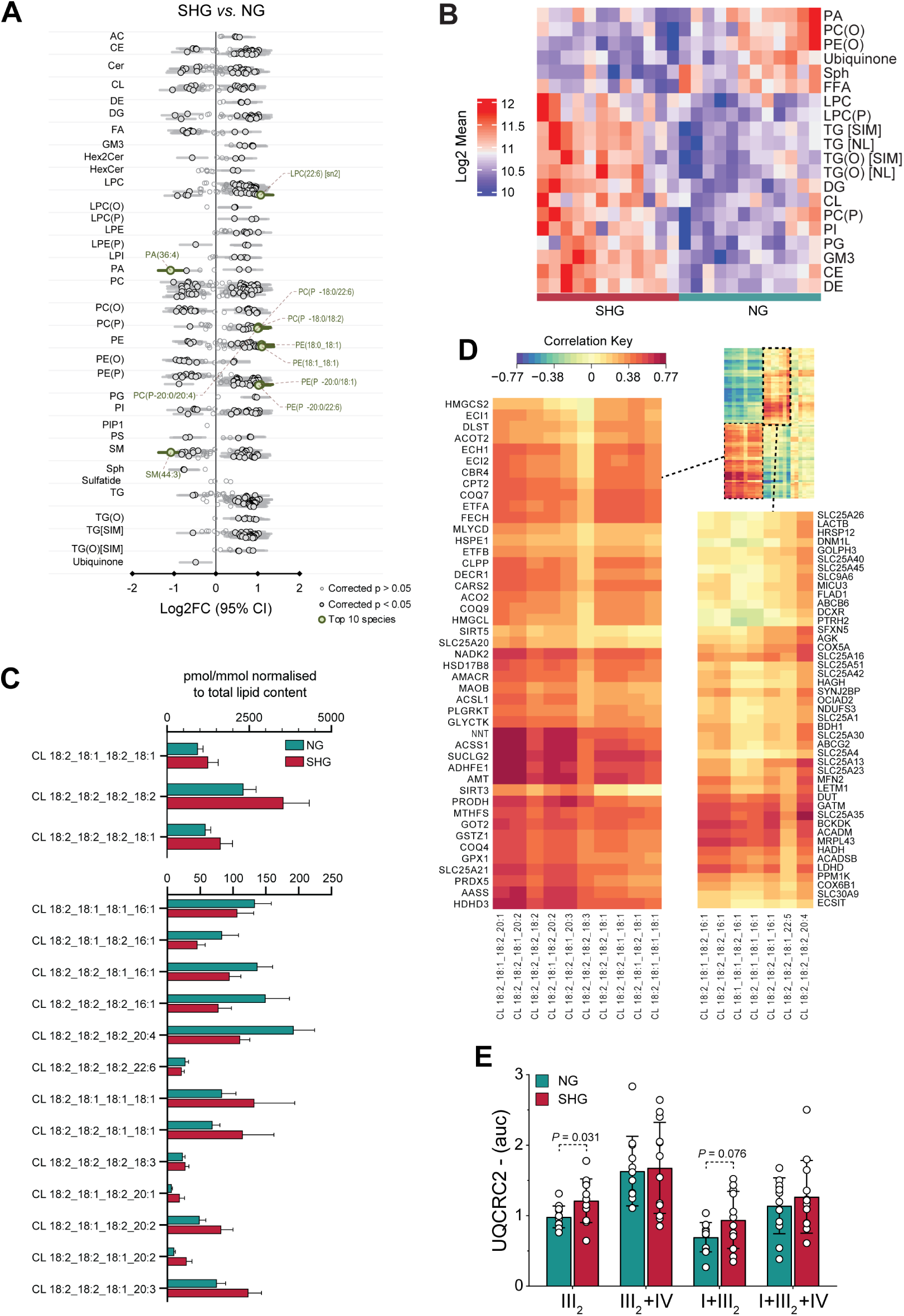
Hyperglycemia induces cardiolipin remodeling. (A) Forrest plot displaying severe hyperglycemia (SHG)-induced changes (expressed as Log_2_FC) in individual lipid species obtained from lipidomic analysis. (B) Hierarchical clustering heat map analysis of lipid classes between SHG and normoglycemia (NG). (C) Bar graph of all differentially expressed cardiolipins (CLs) between SHG and NG in pmol/mmol normalized to total lipid content. Two graphs are displayed separately to ensure optimal visibility due to large differences in the scales. Data are mean ± SD. (D) Regularized canonical correlation analysis between the CL species identified and all the differentially expressed proteins between SHG and NG (normalized log2 absolute values). All analyses ran on mitochondrial fractions isolated from rat kidney cortex; n = 12 (NG) and 11 (SHG). (E) Quantification analysis of supercomplexes from BN-PAGE immunoblots against ubiquinol-cytochrome *c* reductase core protein 2 (UQCRC2); representative images are presented in Figure S5 A and B; n = 12/group. Data are mean ± SD. For panels (C) and (E), outliers were first removed using the ROUT test with Q = 1%; normality was tested with a Shapiro-Wilk test; data were analyzed by one-way ANOVA followed by Tukey post hoc test. Dots represent individual data points.

The biogenesis of the SLC25 family members is dependent on cardiolipins (CLs) (Claypool, 2009). CLs are a unique mitochondrial specific subset of glycerophospholipids found almost exclusively within the IMM, which act as essential structural components in a number of mitochondrial protein complexes and are key for optimal mitochondrial bioenergetics and architecture (Paradies et al., 2014; Prola et al., 2021). Owing to the profound dysregulation of several members of the SLC25 family observed in our proteomics assessment, we investigated the extent of CL remodeling induced by severe hyperglycemia. Our data showed an overall increase in the CL class in SHG vs. NG kidneys (Figure 6B; Table S9). However, in-depth analysis demonstrated extensive remodeling within the CL class, whereby 16 of the 23 identified CL species were differentially expressed in SHG (Figure 6C; Table S9). In particular, we identified an increase in many of the most abundant CLs (e.g., CL 18:2_18:1_18:2_18:1, CL 18:2_18:2_18:2_18:2, CL 18:2_18:2_18:2_18:1), and a decrease in CLs predominately containing 16:1s (Figure 6C; Table S9). To exploit the power of the omics techniques, we performed a regularized canonical correlation analysis between the changes induced by severe hyperglycemia (*vs.* NG) in all differentially expressed proteins and all the identified CLs (Figure 6D). We found a strong positive correlation between i) all 45 upregulated proteins, which were enriched in the TCA cycle and fatty acid β-oxidation metabolism (Figure 4D), and the CLs upregulated by SHG (Figure 6D, large inset); and ii) all 45 downregulated proteins, which were involved in cofactor transport, respiratory ETC, cellular amino acid catabolic process, and mitochondrial transport (Figure 4C and D) and CLs abundant in 16:1 species, which were both decreased in SHG (Figure 6D, small inset). These findings highlight that the specific remodeling of different CLs induced by severe hyperglycemia may be associated with alterations in distinct metabolic and biological processes, consistent with previous findings in different models of experimental diabetes (Ducasa et al., 2019; Sas *et al*., 2021; Tan *et al*., 2020b). Future research should also aim to determine this possible association and if this specificity is also reflected in changes in the overall shape and development of the mitochondrial membranes. Finally, lowering blood glucose to moderate hyperglycemia stabilized the CL lipidome, whereby the CL profile was more consistent with that of normoglycemia (Figure S4A; Table S9). SHG *vs.* MHG differences are also presented in Figure S4B and Table S9.

Studies in yeast demonstrate that CL dysregulation induces mild defects in the assembly and stability of supercomplexes of the ETC (Brandner et al., 2005; Claypool, 2009). Owing to these findings and to the downregulation of OXPHOS complexes induced by SHG, we performed BN-PAGE to investigate if SHG resulted in perturbed supercomplex formation and structure. We observed mild redistribution of the supercomplexes into higher molecular weight assemblies, running slower on the gels, and the destabilization of CIII (Figure 6E; Figure S5A, B, and C). Taken together, these findings suggest that severe hyperglycemia may affect the stability and function of supercomplexes, an effect that may be induced by CL remodeling.

### Correlations between markers of DKD and differentially expressed proteins or metabolites highlight new potential markers of the disease

To identify new potential protein markers of DKD, we performed a correlation analysis between all the differentially expressed proteins induced by SHG *vs.* NG and markers of renal function or injury (i.e., plasma cystatin C [Figure S6] and urinary albumin [Figure S7]). Our analysis identified five proteins that were significantly correlated with plasma cystatin C (Figure 7A). SLC25A30, also known as kidney mitochondrial carrier protein 1 (KMCP1), was positively associated with plasma cystatin C. It is a kidney-specific protein activated when mitochondrial metabolism is increased, in particular when the cellular redox balance tends toward a pro-oxidant status (Haguenauer *et al*., 2005). HSD17B8, a protein involved in fatty acid biosynthesis (Venkatesan et al., 2014), and BDH1 were also correlated with cystatin C (Figure 7A). The strong correlation between cystatin C and BDH1 reaffirms the importance of ketone body metabolism in DKD and the role of ketones as key substrates in fueling the kidney’s energetic demands in DKD. AMT, which catalyzes the degradation of glycine, was negatively associated with cystatin C. There was a negative association between SUCLG2 and plasma cystatin C. SUCLG2 is a subunit of Succinyl-CoA synthetase which catalyzes the reversible reaction involving the formation of succinyl-CoA and succinate.

**Figure 7.**
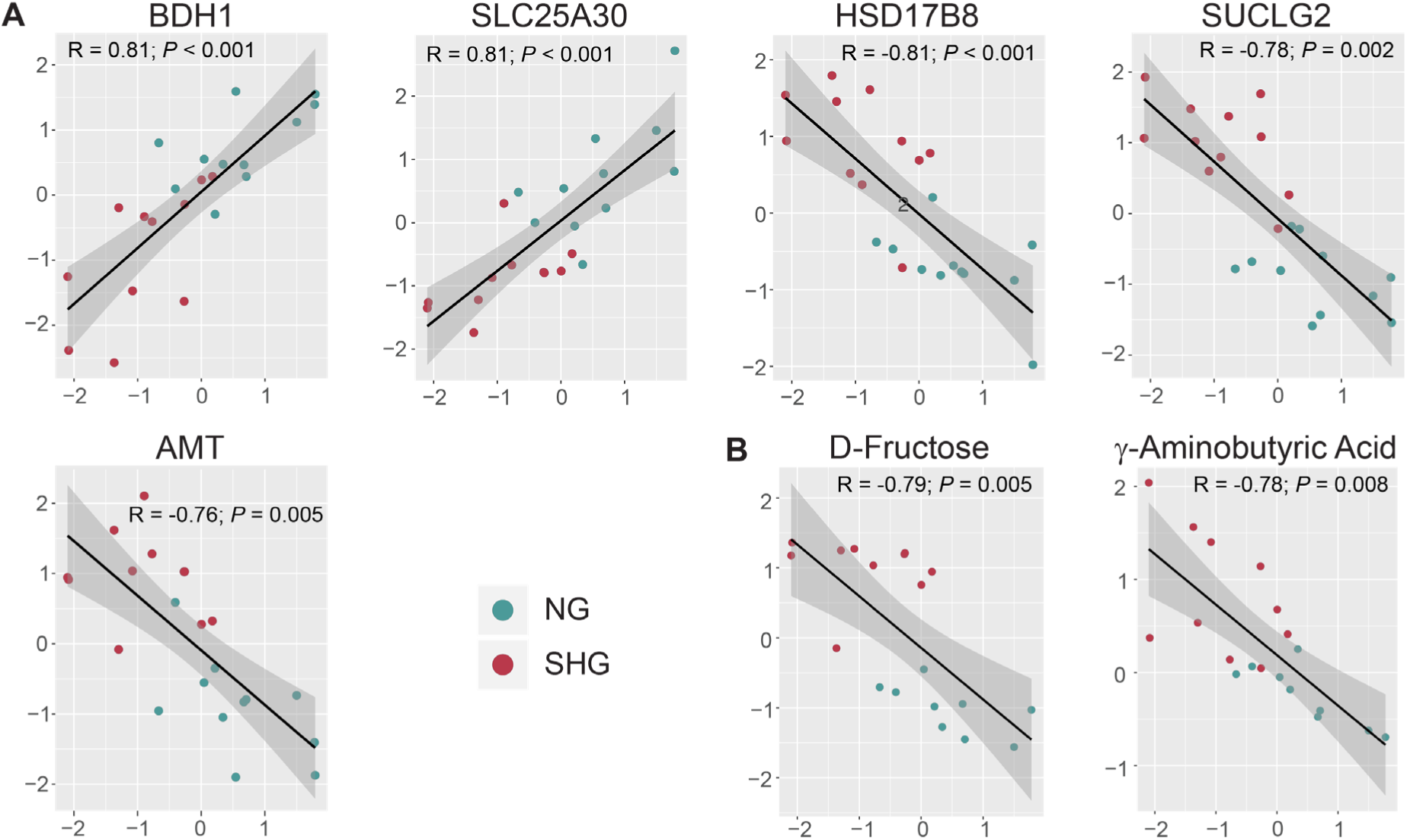
Correlation plots between differentially expressed proteins or metabolites in SHG *vs.* NG and surrogate markers of kidney function (cystatin C). Correlation plots displaying significant correlations between A) differentially expressed proteins or B) differentially expressed metabolites in SHG *vs.* NG and markers of DKD (plasma cystatin C and urinary albumin). Markers values (x-axis) underwent a log2 transformation followed by scaling and were correlated against normalized log2 transformed intensity values for proteins and metabolites (y-axis). Correlation coefficients were calculated using the Spearman method with Holm corrections to account for multiple comparisons; significance was set at *P* < 0.01. Dots represent individual values. n = 10 (NG) and 11 (SHG). The correlations for all differentially expressed proteins or metabolites in SHG *vs.* NG and markers of DKD, regardless of significance, are shown in Figure S6 to S9.

None of the differentially expressed proteins by SHG (*vs.* NG) were significantly correlated with urinary albumin (Figure S7). While cystatin C is a surrogate marker of glomerular filtration, with hyperfiltration generally exhibited in early DKD (as evidenced in the current study by a reduction in plasma cystatin C), albuminuria generally takes place later in the progression of DKD (Sudamrao Garud and Anant Kulkarni, 2014). Therefore, the significant correlations of differentially expressed mitochondria-associated proteins observed with plasma cystatin C, but not with urine albumin, suggest that these mitochondrial proteins may be linked with early dysregulation of kidney function.

We next performed correlation analyses between all the differentially expressed metabolites induced by SHG (*vs.* NG) and plasma cystatin C (Figure S8) or urinary albumin (Figure S9). Both D-fructose and γ-aminobutyric acid were negatively correlated with cystatin C (Figure 7B). The presence of increased fructose levels in SHG kidneys may relate to findings demonstrating endogenous renal fructose production in DKD, which leads to increased uric acid production, as well as kidney inflammation and fibrosis (Nakagawa et al., 2020). In addition, we identified 5 differentially expressed proteins and 19 metabolites that were significantly correlated with HbA_1c_ (Figure S10 and S11, respectively).

## Discussion

This study has shown that titrating blood glucose concentrations *in vivo* directly impacts mitochondrial morphology and bioenergetics as well as intensely remodeling the mitochondrial proteome and the lipidome in the kidney in early DKD. Mitoproteomic analysis revealed profound metabolic disturbances induced by severe hyperglycemia, including upregulation of enzymes involved in the TCA cycle and fatty acid metabolism and enhanced ketogenesis. Untargeted metabolomics and lipidomics confirmed the enrichment of TCA cycle metabolites, an increase in triglyceride concentrations, and extensive and specific cardiolipin remodeling. Lowering blood glucose to moderate hyperglycemia stabilized all three omic landscapes, partially prevented changes in mitochondrial morphology and bioenergetics, and improved kidney injury, with improvement of kidney function (cystatin c), glomerulosclerosis, tubular injury (Kim-1) and markers of fibrosis (TGFβ1).

Taken together with the impairments seen in mitochondrial complex I activity, our data indicate that severe hyperglycemia promotes a metabolic switch towards a more glycolytic phenotype (Warburg effect), a previously described feature of DKD (Zhang *et al*., 2018). In situations of limited oxygen supply, such as the proximal tubule in DKD, where the cell is facing hypoxia, it may switch from mitochondrial respiration, towards increased glycolysis in order to maintain ATP (Hesp et al., 2020).

Data from our exploratory multi-omic workflow demonstrate the importance of mitochondria-associated lipids in the kidney and how they change in relation to hyperglycemia in the setting of diabetes. Collectively, our datasets indicate that altering blood glucose concentration *in vivo* changes lipid uptake and metabolism in the kidney as shown by a profound remodeling of the lipidome, and based on the mitoproteomics findings, dysregulation of lipid metabolism. Indeed, there has been increasing attention on the role of kidney lipid metabolism in DKD, with studies indicating that there may be defects in lipid uptake into the mitochondria, increased lipid biosynthesis, but also defective fatty acid oxidation (Stadler *et al*., 2015). It has been suggested that lipid accumulation in the kidney drives lipotoxicity resulting in fibrosis (Nicholson et al., 2020; Stadler *et al*., 2015), though the precise molecular mechanisms are not known. Although an increase in triglycerides induced by severe hyperglycemia was one of the predominant features of our lipidomics analysis, it is likely that functional changes to specific lipid classes, not just lipid abundance, are important in the development of DKD. In particular, our lipidomics output has shown remodeling of cardiolipins with increasing glucose concentrations. Severe hyperglycemia also induces upregulation of the carnitine shuttle and other enzymes involved in transport of fatty acids to mitochondria for β-oxidation. The same milieu upregulates enzymes involved in the β-oxidation of unsaturated fatty acids, and decreases branched-chain amino acid and fatty acid catabolism. In addition, related to lipid metabolism, is the alteration found in ketone body metabolism and increased ketogenesis. However, our study was not able to investigate *de novo* lipogenesis in the kidney and this remains as yet, a relatively neglected area of study.

In conclusion, using a multi-omic approach, we show that blood glucose trajectories are a key determinant of adaptive changes in mitochondrial flexibility, signaling and fatty acid utilization in the kidney in early diabetes. This study provides insight into mitochondrial function and energy generation in the kidney during severe hyperglycemia and has implications for therapeutic strategies aimed at the reinvigoration of mitochondrial function and signaling in diabetes.

## Methods

### Experimental Animal Model

Male Sprague Dawley rats were housed in groups of three rats per cage in a temperature-controlled environment, with a 12 h light/dark cycle and *ad libitum* access to food and water. Experimental diabetes was induced in six week old male Sprague Dawley rats (200-250 g, n = 35) by *i.v*. injection of streptozotocin (55 mg/kg, sodium citrate buffer pH 4.5) following an overnight fast, as previously described (Rosenkranz et al., 2003). One group of rats received citrate buffer vehicle (0.42% in sterile saline, pH 4.5) as a non-diabetic control with normal blood glucose (NG) (n = 16). One week following STZ treatment, diabetic rats were further assigned to two groups: standard insulin therapy (n = 17 rats), resulting in severe hyperglycemia (SHG) and intensive insulin therapy (n = 19 rats), resulting in moderate hyperglycemia (MHG) using a single daily insulin injection (long-lasting Humulin NPH; Eli Lilly, Indianapolis, USA) to titrate blood glucose levels to >28 mM (1-2 units, s.c. per day) and ∼20 mM (6-7 units, s.c. per day) as required, respectively. Blood glucose and body weight were monitored weekly. Blood glucose was measured using a handheld glucometer (Accutrend; Boehringer Manheim Biochemica, Manheim, Germany) during the study time course. After the completion of the study, plasma glucose was measured using a colorimetric glucose assay kit from Cayman Chemical Company (Ann Arbor, MI, USA), performed according to the manufacturer’s instructions. Hemoglobin A_1c_ (HbA_1c_) was determined by a Cobas Integra 400 autoanalyzer (Roche Diagnostics Corporation, USA). Plasma C-peptide was determined using a commercially available ELISA kit (Alpco, Salem, NH, USA) according to the manufacturer’s instructions. In the final week of the study, rats were placed individually into metabolic cages (Iffa Credo, L’Arbresele, France) for 24 hours to collect urine. Eight weeks after STZ-induced diabetes, rats were euthanized with an intraperitoneal injection of ketamine-xylazine (100 and 12 mg/kg, respectively) and the kidneys were rapidly dissected, weighed, and snap-frozen or placed in 10% neutral buffered formalin (v/v) for fixation before paraffin embedding. In a subset of rats (*n* = 3-4 per group), renal cortices were processed for transmission electron microscopy (TEM) as described below.

### Study approval

All animal experiments complied with the National Health and Medical Research Council (NHMRC) of Australia code of practice for the care and use of animals for scientific purposes and all procedures were approved by the Alfred Medical Research Educational Precinct (AMREP) Animal Ethics Committee (study approval number E/1519/2014/B).

### Renal Function and Morphometry

ELISAs were used to measure urinary albumin excretion (Bethyl Laboratories, Montgomery, TX, USA), plasma cystatin C (BioVendor, Mordice, Czech Republic) and Kidney Injury Molecule-1 (Kim-1, Abcam, Cambridge, United Kingdom) as per the manufacturer’s instructions. Glomerulosclerotic index (GSI) was assessed in a blinded fashion in Periodic Acid Schiff (PAS)-stained sections as previously described (Saito et al., 1987). Biologically active TGF-β1 was measured in urine using the TGF-β1 Emax Immunoassay System (Promega, Madison, WI) according to the instructions. Prior to measurement, samples were acid activated with 1M HCl followed by neutralization to pH 7.0 with 1M NaOH. Values were expressed as pg/mg of protein as determined by the bicinchoninic acid (BCA) method (Pierce, Rockford, IL, USA). Paraffin sections of kidneys were used to stain for collagen type IV using a goat polyclonal collagen IV antibody (Southern Biotech, Birmingham, AL, USA) (Thallas-Bonke et al., 2008).

### Mitochondrial isolation for assessment of enzymatic activity, ROS generation, and BN-PAGE

Mitochondria were isolated by differential centrifugation. Freshly harvested renal cortex (50 mg) was finely minced and gently homogenized with glass Teflon tissue grinders in 2 ml ice-cold isolation medium, pH 7.2 (70 mM sucrose, 210 mM mannitol, 5 mM HEPES, 1mM EGTA). The homogenate was centrifuged at 800 g for 5 min at 4°C and the resulting supernatant was centrifuged at 8,000 g for 10 min at 4°C. After washing with 0.5 ml ice-cold isolation buffer, the mitochondrial pellet was resuspended in 200 μl ice-cold isolation medium. Total protein was determined by the bicinchoninic acid method according to the manufacturer’s instructions (BCA Protein Assay Kit, Pierce-Thermo Fisher Scientific, Melbourne, Australia).

### Citrate Synthase activity

Mitochondrial citrate synthase (CS) activity was performed as previously described (Granata et al., 2016) with minor modifications. This assay measures the rate of citrate synthase activity by linking the release of Coenzyme A to the colorimetric agent DTNB 5,5-dithiobis-2-nitrobenzoate (Ellman’s reagent). Ten μg of previously sonicated isolated mitochondria was added in triplicate to a microtiter plate in the presence of the following reagents: 500 μM acetyl coenzyme A, 100 μM 5,5’-dithiobis(2-nitrobenzoic acid) and 100 mM Tris buffer (pH 8.3) at 25°C. After addition of 15 μl of 10mM oxaloacetic acid (final concentration 600 μM), the plate was immediately placed in a spectrophotometer at 25°C (Clariostar Plus, BMG Labtech, Mornington, VIC, Aus), and after 15 s of linear agitation, absorbance at 412 nm was recorded every 10 s for 5 min. The change in absorbance per minute was calculated within the linear range of the plot (∼1-2 min). After correction for the blank, citrate synthase activity was calculated using the molar extinction coefficient of TNB at 412 nm of 13.6 mM^-1^ cm^-1^, and was standardized to total protein.

### Complex I activity

Respiratory chain complex I was determined as previously described (Van Bergen et al., 2014) in mitochondria isolated from renal cortex as described above. Briefly, samples were assayed in triplicate in a Corning® UV-Transparent 96-well plate (Sigma-Aldrich Pty Ltd, Sydney, Australia) after adding 8 μg of previously sonicated mitochondrial isolate to a reaction mixture containing: 250 mM sucrose, 50 mM Tris, 1 mM EDTA, 50 μM decylubiquinone, 2 mM KCN, 50 μM Nicotinamide adenine dinucleotide (NADH) at pH 7.4. The oxidation of NADH was then monitored in a spectrophotometer (Clariostar Plus, BMG Labtech, Mornington, VIC, Aus) at 340 nm for 3 min at 25°C. The linear phase of the reaction (∼1 min) was used for the determination of maximal complex I activity. A parallel assay was performed (in triplicate) in the presence of 4 μg/ml rotenone (an inhibitor of complex I) to determine the rate of rotenone-insensitive activity, and this value was subtracted from the maximal activity value obtained in the absence of rotenone. To account for variations in Mito_VD_ between samples, complex I activity was expressed relative to citrate synthase activity (Frazier et al., 2020).

### Hydrogen peroxide production

Hydrogen peroxide production in mitochondria isolated from the renal cortex was measured using a commercially available Amplex Red H_2_O_2_ assay kit (Invitrogen), as previously described (Forbes et al., 2013). In brief, 16 µl of either 10 mM glutamate and 10 mM malate (G/M), 10 mM succinate and 2.5 µM rotenone (S/R), or Krebs-Ringer phosphate glucose solution (No Substrate; NS) were added to a microplate well containing 20 µl of mitochondrial isolate, in duplicate for each substrate mixture. 100 µl of a pre-warmed reaction solution, containing 50 μM Amplex Red reagent and 0.1 U/ml horseradish peroxidase, was added to each well and the plate was incubated for 30 minutes at 37°C. Following incubation, florescence was measured using a fluorescence spectrophotometer (Clariostar, BMG Labtech) set to excitation 540 nm, emission 590 nm. Data are expressed as relative to no substrate control.

### Transmission electron microscopy and analysis of mitochondrial morphology

Immediately following exsanguination, renal cortices were fixed in Karnovsky’s fixative, 2% paraformaldehyde, 2.5% glutaraldehyde in 0.1M cacodylate buffer (pH 7.2) for four hours at room temperature. The samples were then rinsed three times in fresh buffer for 15 minutes each before post-fixing in 1% Osmium tetroxide, 1.5% potassium ferrocyanide in 0.1M cacodylate buffer for 2 hours at room temperature. Fixed tissue samples were rinsed three times in milliQ H_2_O for ten minutes each, before being dehydrated in increasing concentrations of ethanol followed by acetone. Following dehydration, the tissues were infiltrated with increasing concentrations of epon-araldyte resin in acetone. After a second change of 100 % resin the samples were embedded in fresh resin in flat bottom BEEM capsules and the resin was allowed to polymerize in an oven at 65 °C for 48 hours. The embedded tissues in resin blocks were sectioned with a diamond knife on a Leica Ultracut S microtome and ultra-thin sections (90 nm) were collected onto formvar-coated 100 mesh hexagonal copper grids. The sections on grids were sequentially stained with 1% uranyl acetate for 10 mins and Triple Lead Stain for 5 min (Sato, 1968) and viewed in a Jeol JEM-1400Plus transmission electron microscope equipped with a Matataki flash camera at 80keV. Five to fifteen images of proximal tubule epithelial cells per kidney section were randomly collected for each rat (*n* = 3-4 rats per group) at x15,000 magnification. Measurements of mitochondrial morphology (form factor, aspect ratio, circularity, and roundness) within a given image were measured using ImageJ (National Institutes of Health, Bethesda, MD, USA). Measurements of Mito_VD_ were obtained by normalizing the total mitochondrial area by the subcellular area subtracted from the nuclear area, within a given image.

### BN-PAGE

Mitochondrial isolates (15 μg) were separated by electrophoresis using 3-12% NativePAGE gels (Life Technologies Australia, Mulgrave, Australia) to assess supercomplex formation using a 4 g/g digitonin/protein ratio, as previously described (Granata et al., 2021a). The following primary antibodies were used: NDUFA9 (ab14713) and UQCRC2 (ab14745) (both Abcam, Cambridge, MA, USA). Protein bands were visualized using a Bio-Rad ChemiDoc imaging system and were quantified using Bio-Rad Image Lab 5.0 software (Bio-Rad laboratories, Gladesville, NSW, Australia). An internal standard (consisting of a mix of all samples analyzed) was loaded in each gel, so that each lane could be normalized by this value, to reduce gel-to-gel variability.

### Mitochondrial isolation for mitochondrial proteomic and lipidomics analysis

Approximately 30 mg of frozen renal cortex were homogenized in 3 ml ice-cold isolation medium, pH 7.6 (20 mM HEPES-KOH pH 7.6, 220 mM mannitol, 70 mM sucrose, 1 mM EDTA, and 0.5 mM phenylmethylsulfonyl fluoride [PMSF]) at 4°C with 40 strokes of a Teflon on glass Potter-Elvehjem homogenizer fitted to a drill rotating at ∼ 800 rpm. Homogenates were centrifuged at 1,000 g for 10 min at 4°C and the supernatant was further centrifuged at 12,000 g for 15 min at 4 °C. The pellet was resuspended in 200 μL of sucrose buffer (10 mM HEPES-KOH pH 7.6, 0.5 M sucrose) and a protein concentration assay was performed (BCA Protein Assay Kit, Pierce-Thermo Fisher Scientific, Melbourne, Australia). 100 μg of isolated mitochondria were collected and centrifuged at 12,000 g for 15 min at 4 °C and after removal of the supernatant the pellet was frozen at −80 °C for subsequent proteomics analysis.

### Proteomics analysis

100 μg of mitochondrial isolate were solubilized in 20 μL of 8 M urea, 40mM chloroacetamide (CAA), 10 mM tris(2-carboxyethy)phosphine (TCEP), 100 mM Tris pH and sonicated in a water bath for 15 min. After incubation at 37°C with shaking (1500 rpm), 60 μL of MilliQ H_2_O was added to dilute the urea to 2 M, after which samples were digested overnight at 37 °C with trypsin (Promega, Fitchburg, WI,) in a 1:60 trypsin to protein ratio. Samples were next acidified with 10% trifluoro acetic acid (TFA) and subsequently centrifuged at 20,100 g at RT. The supernatant was loaded onto equilibrated stage tips prepared in house by stacking 3×14G styrene divinylbenzene (SDB)-XC layers (Empore Solid Phase Extraction Products, 3M, Eagan, MN, USA) into a 200 μL Eppendorf tip (Granata et al., 2021b; Kulak et al., 2014). Stage tips were washed with 100 μL 0.1% TFA, 2% acetonitrile (ACN) by centrifugation at 2,300 g at RT and peptides were eluted with 100 μL 0.1% TFA, 80% ACN by centrifugation at 2,300 g at RT. Peptides were vacuum dried (SpeedVac Vacuum Concentrator, Thermo Fisher Scientific, Melbourne, Australia) and reconstituted in 25 μL 0.1% TFA, 2% ACN, vortexed thoroughly, and sonicated for 15 min before being centrifuged at 20,100 g for 5 min at RT. The supernatant was collected, peptide concentration was estimated via Direct Detect® Infrared Spectrometer (Merck, Bayswater, VIC, Australia), and samples were transferred to HPLC vials for proteomics analysis.

Approximately 800 ng of peptides were analyzed sequentially by online nano-HPLC/electrospray ionization-MS/MS on a Q Exactive Plus connected to an Ultimate 3000 HPLC (Thermo Fisher Scientific, Melbourne, Australia) as previously described (Granata *et al*., 2021a). Briefly, peptides were initially loaded onto a trap column (Acclaim C18 PepMap nano Trap × 2 cm, 100 μm I.D, 5-μ m particle size and 300-Å pore size; Thermo Fisher Scientific) at 15 μl min^−1^ for 3 min and subsequently switched to the analytical column (Acclaim RSLC C18 PepMap Acclaim RSLC nanocolumn 75 μm × 50 cm, PepMap100 C18, 3-μm particle size 100-Å pore size; Thermo Fisher Scientific). Peptide separation was carried out at 250 nL min^−1^ using a nonlinear ACN gradient of buffer A (2% ACN, 0.1% FA) and buffer B (80% ACN, 0.1% FA), from 2.5% to 35.4% buffer B, followed by ramp to 99% buffer B over 120 min (155 min total acquisition time to accommodate for void and equilibration volumes). Data were collected in Data Dependent Acquisition (DDA) positive mode with a scan range of 375–1800 m/z in MS1, and higher energy collisional dissociation (HCD) was used for MS/MS scans of the 12 most intense ions with z ≥ 2. Other instrument parameters were: MS1 scan resolution: 70,000 (at 200 m/z); MS maximum injection time: 50 ms; AGC target: 3E+6; normalized collision energy: 27%; Isolation window: 1.8 Da; MS/MS resolution: 17,500; MS/MS AGC target: 1E+5; MS/MS maximum injection time: 100 ms; minimum intensity: 1E+3; dynamic exclusion: 15 s.

Analysis of raw mass spec files was carried out using the MaxQuant (version 1.6.1.0) software (Cox and Mann, 2008) searching against the Uniprot *Rattus norvegicus* database containing reviewed, canonical and isoform variants in FASTA format (June 2018) and a database containing common contaminants by the Andromeda search engine (Cox et al., 2011). Data analysis was performed with variable (methionine oxidation [M] and N-terminal acetylation) and fixed (cysteine carbamidomethylation [C]) modifications with label free quantitation, and matching between runs. A false discovery rate (FDR) of *P* < 0.01 for proteins and peptides was applied by searching a reverse database. The ‘match from and to’ and ‘match between runs’ options were enabled with a 2-min match time window. Protein quantification was performed on a minimum ratio count of 2 unique or razor peptides.

### Bioinformatic analysis of proteomics data

For bioinformatic analysis of the proteomic dataset, the R package *limma* (Ritchie et al., 2015) was used to conduct the differential expression analysis of MaxQuant LFQ intensities after first performing normalization by cyclic loess within the *limma* package. Identifications labelled by MaxQuant as ‘only identified by site’, ‘reverse’ and ‘potential contaminant’ were removed. Proteins with less than 70% valid values were removed and remaining missing data was imputed using nearest neighbor averaging with a k=4 from the *impute.knn* package in R (Hastie et al., 2021). To identify the mitochondrial proteins, all “Known Mitochondrial” proteins identified using the IMPI database (Smith and Robinson, 2019) were subset from the rest of the dataset and used for subsequent analysis. Differential expression analysis was performed between pairs of groups. Linear modelling was determined using eBayes in the *limma* package. The resulting differential expression values were filtered for an adjusted *P* value < 0.05 using the Benjamini-Hochberg method. Heatmaps were produced using unsupervised hierarchical clustering using the “ward.D2” method. Gene ontology of the clusters was determined by taking the proteins identified in the cluster and performing an enrichment analysis using the ClueGO (v2.5.6) application in Cytoscape (v3.7.1) using default settings except for a GO tree interval of 3 to 5 and a network specificity of medium, with the Biological Processes Ontology and Reactome Ontology switched on. Volcano plots were created using the *EnhancedVolcano* package in R (Blighe et al., 2021). A complete list of all the identified proteins and post-normalization differentially expressed proteins between group pairs, in their respective clusters and their annotations, are presented in Table S1 to S3; enrichment analysis is presented in Table S4.

### Kidney cortex metabolite extraction and sample derivatization for metabolomics analysis

Metabolite extraction was carried out as previously described (Bachem et al., 2019). Briefly, ∼20-30 mg of kidney cortex was homogenized under cryogenic conditions in cryomill tubes containing beads for homogenization (Precellys bead-mill with a Cryolys attachment, Bertin Technologies, France) and 600 mL of 3:1 methanol:water (v/v) containing 0.5 nmol 13C6-sorbitol and 5 nmol 13C5,15N-valine as internal standards. Homogenates (480 mL) were subsequently vortexed in fresh Eppendorf tubes containing 120 ml chloroform. The resultant extracts were centrifuged to pellet cell debris and precipitated protein. The supernatant was used for subsequent analysis. In addition, an aliquot from each sample was pooled and re-aliquoted to generate pooled biological quality controls (PBQC). Samples and PBQCs were evaporated dry by speed vacuum centrifugation and then derivatized online using the Shimadzu AOC6000 autosampler robot with methoxyamine hydrochloride (30 mg/mL in pyridine) and N, O - bis (trimethylsilyl) trifluoroacetamide [BSTFA] + 1% chlorotrimethylsilane [TMCS] (both Thermo Fisher Scientific, Waltham, USA). Samples were left for 1 h before 1 µL was injected onto the GC column using a hot needle technique. Split (1:10) injections were performed for each sample.

### Metabolomics analysis

The GC-MS system consisted of an AOC6000 autosampler, a 2030 Shimadzu gas chromatograph and a TQ8040 quadrupole mass spectrometer (Shimadzu, Japan), which was tuned according to the manufacturer’s recommendations using tris-(perfluorobutyl)-amine (CF43). GC-MS was performed on a 30 m Agilent DB-5 column with 1 µm film thickness and 0.25 mm internal diameter. The injection temperature (Inlet) and the MS transfer line were both set at 280°C and the ion source adjusted to 200°C. Helium was used as the carrier gas at a flow rate of 1 mL/min and argon gas was used as the collision cell gas to generate the multiple reaction monitoring (MRM) product ion. The analysis was performed under the following temperature program; start at injection 100°C, a hold for 4 minutes followed by a 10°C min^-1^ oven temperature ramp to 320°C followed by a final hold off of 11 minutes. Approximately 520 quantifying MRM targets were collected using Shimadzu Smart Database along with qualifier for each target, which covers about 350 endogenous metabolites and multiple ^13^C labelled internal standards. Both chromatograms and MRMs were evaluated using the Shimadzu GCMS browser and LabSolutions Insight software.

### Bioinformatic analysis of metabolomics data

For bioinformatic analysis of the metabolomic dataset, pareto scaling was applied to the logged concentrations of the metabolites for normalization. Differential expression analysis was completed similarly to the proteomics analysis between all group pairs. Linear modelling was determined using eBayes in the *limma* package (Ritchie *et al*., 2015). The resulting differential expression values were filtered for an adjusted *P* value < 0.05 using the Benjamini-Hochberg method. Heatmaps were produced using unsupervised hierarchical clustering using the “average” method. To highlight the correlation between the differentially expressed proteins and metabolites, a regularized canonical correlation analysis (rCCA) was performed using the MixOmics package in R (Rohart et al., 2017) which yields linear combinations of the variables from each original dataset in order to explore the associated canonical correlation between the two components. The raw, normalized and differentially expressed metabolomics data are presented in Tables S5 and S6. An over representation analysis for the metabolites was performed using the online platform MetaboAnalyst (Pang et al., 2021); the differentially upregulated and downregulated metabolites were separately entered into MetaboAnalyst with the top five significant terms from the Small Molecular Pathway Database (SMPDB) shown in Figure 5B. All the terms identified by MetaboAnalyst are presented in Table S7.

### Lipidomic analysis

Lipids were extracted from mitochondrial isolates using a single-phase chloroform/methanol extraction as described previously (Granata *et al*., 2021a). Briefly, 20 volumes of chloroform:methanol (2:1) were added to the sample along with a series of internal standards. Samples were vortexed and centrifuged on a rotary mixer for 10 min. Following 30 min of sonication in a sonicator bath, samples were rested for 20 min before being centrifuged at 13,000 g for 10 min. Supernatants were transferred into a 96 well plate, dried down, and reconstituted in 50 µL H_2_O saturated butanol, before being sonicated for 10 min. Following the addition of 50 µL of methanol with 10 mM ammonium formate, samples were centrifuged at 4000 rpm on a plate centrifuge and transferred into glass vials with inserts for mass spectrometry analysis.

### Targeted lipidomics analysis

LC-MS/MS was performed according to previously published methods, with slight modifications (Huynh et al., 2019). Sample extracts were analyzed using either (i) an AB Sciex Qtrap 4000 mass spectrometer coupled to an Agilent 1200 HPLC system for CL assessment, as described previously (Granata *et al*., 2021a) or (ii) an Agilent 6490 QQQ mass spectrometer coupled with an Agilent 1290 series HPLC system for assessment of all other lipid species (Huynh *et al*., 2019). Lipids run on the Agilent 6490 were measured using scheduled multiple reaction monitoring with the following conditions: isolation widths for Q1 and Q3 were set to “unit” resolution (0.7 amu), gas temperature 150°C, nebulizer 20 psi, sheath gas temperature 200°C, gas flow rate 17 L/min, capillary voltage 3500 V and sheath gas flow 10 L/min. The list of MRMs used and the chromatographic conditions were described previously (Huynh *et al*., 2019).

### Bioinformatic analysis of lipidomics data

Lipid classes were log transformed before being normalized using the NormaliseBetweenArrays function in *Limma* (Ritchie *et al*., 2015). Differential expression analysis was completed similarly to the proteomics analysis between all pairs of groups. Linear modelling was determined using eBayes in the *limma* package (Ritchie *et al*., 2015). The resulting differential expression values were filtered for an adjusted *P* value < 0.05 using the Benjamini-Hochberg method. Forest plots were generated using LogFC results from *Limma* analysis. Heatmaps were produced using unsupervised hierarchical clustering using the “ward.D2” method. Lipid species were analyzed in the same way as the classes. Changes in the overall normalized pmol/mmol concentration of the differentially expressed CLs between SHG and NG is displayed in Figure 6C. To highlight the correlation between all the CLs and differentially expressed proteins, a rCCA was performed using the MixOmics package in R (Rohart *et al*., 2017). A complete list of all the identified lipid species and classes and post-normalization differentially expressed lipid species and classes between group pairs are presented in Table S8 and S9.

### Correlation analyses

Correlation analyses between the differentially expressed proteins or metabolites and markers for DKD (cystatin C and albumin) or diabetes (HbA_1c_) were conducted in R. Values of markers underwent a log2 transformation followed by scaling and were correlated with normalized log2 transformed intensity values for protein or concentrations for metabolites. Correlation coefficients were calculated using the Spearman method with Holm corrections to account for multiple comparisons. The level of statistical significance for all correlations was set at *P* < 0.01.

### Statistical analysis

All values are reported as means ± SEM, unless otherwise specified. For non-omics analyses outliers were first removed using the ROUT method set at Q = 1% (Motulsky and Brown, 2006). Normally distributed datasets (Shapiro-Wilk test *P* > 0.05) were analyzed by one-way ANOVA followed by Tukey’s post hoc testing when comparing 3 groups, or by unpaired t-test when comparing 2 groups. Non-normally distributed datasets (Shapiro-Wilk test *P* < 0.05) were analyzed using the non-parametric Kruskal-Wallis test followed by Dunn’s post hoc testing when comparing 3 groups, or by the non-parametric Mann-Whitney test when comparing 2 groups. The level of statistical significance was set at *P* < 0.05. GraphPad Prism v. 8.4.2 (GraphPad Software, San Diego, California, USA) was used for all statistical non-omics analyses.

### Data Availability

Source data to interpret, verify and extend this research is provided with this paper. The mass spectrometry proteomics data has been deposited in the ProteomeXchange Consortium via the PRIDE partner repository under accession code PXD037705. The lipidomics and metabolomics data have been deposited in the NIH Common Fund’s National Metabolomics Data Repository (NMDR) website, the Metabolomics Workbench, using DataTrack ID 3561, and will be available upon acceptance of the manuscript. The R scripts used for all omics analyses described above are deposited on Zenodo and available through <u>DOI: 10.5281/zenodo.7246478</u>. There are no restrictions placed on accessibility of this code.

### Materials availability

This study did not generate new unique reagents.

## Acknowledgments

The authors would like to thank the following people for their technical input: Maryann Arnstein, Edy Xie, Natalie Kozlov, Gavin Higgins and Runa Lindblom. We would like to thank Joan Clark, Adam Costin and Ben Padman from the Ramaciotti Centre for Cryo Electron Microscopy, Monash University, Australia for assistance with electron microscopy. The authors acknowledge Servier Medical Art (Creative Commons License). Illustrations were created with BioRender.com. This work was completed with support from the National Health and Medical Research Council of Australia (NHMRC) via NHMRC Fellowships (1140851 and 2009732 to DAS; 1197190 to KH, 1059960 to RHR) and grants (1140906 to DAS; 1101309 to MTC, 1045140 to RHR). MTC is the recipient of a Career Development Award from JDRF Australia (4-CDA-2018-613-M-B), the recipient of the Australian Research Council Special Research Initiative in Type 1 Juvenile Diabetes. We acknowledge the Ramaciotti Centre for Cryo Electron Microscopy, Monash University, a node of Microscopy Australia. In addition, we acknowledge the support of the Mito Foundation for the provision of instrumentation and the assistance of the Bio21 Melbourne Mass-Spectrometry and Proteomics Facility.

## Author contribution statement

RR and MTC conceived the study; CXQ, JA and EJ ran the animal study; VTB, CG, AL, MS, GR and DS collected the data. CG, VTB, NJC, KH, GR, DS and MTC researched and analyzed the data; CG, MTC and DS wrote the paper. All authors revised and critically discussed the manuscript. All persons designated as authors qualify for authorship, and all those qualifying for authorship are listed. All authors have read and approved the final manuscript.

## Competing interest

The authors declare no conflict of interest.

## Inclusion and Diversity Statement

We support inclusive, diverse, and equitable conduct of research

## Lead contact statement

Further information and requests for resources and reagents should be directed to and will be fulfilled by the lead contact Melinda Coughlan

## Notes

### Competing Interest Statement

The authors have declared no competing interest.

