## Supplemental Figures for "Deep multi-omic profiling reveals extensive mitochondrial remodeling driven by glycemia in early diabetic kidney disease"

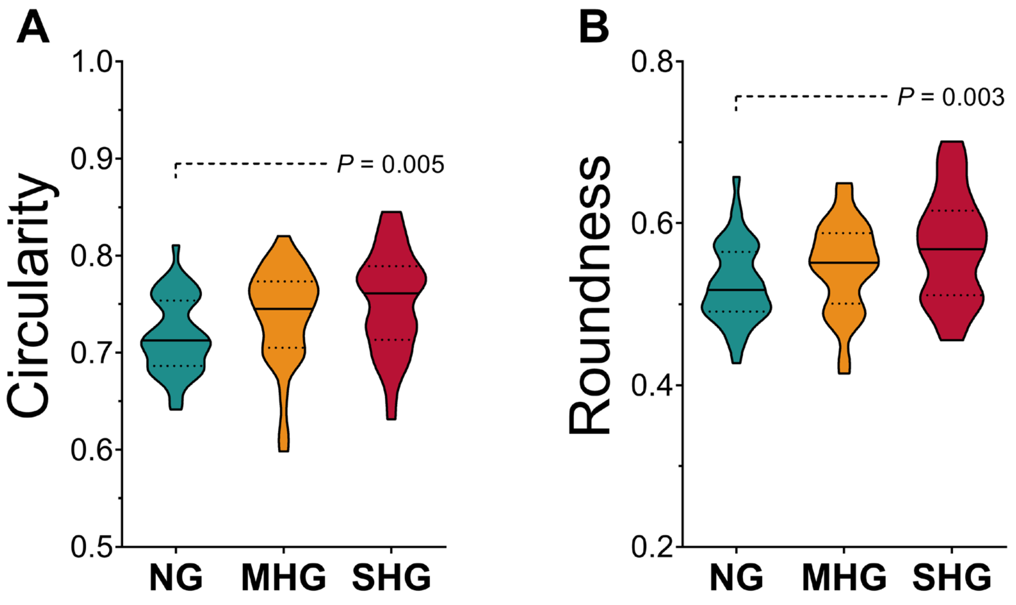


**Figure S1. Mitochondrial morphology.** (A) Mitochondrial circularity; (B) mitochondrial roundness, determined from electron micrographs of renal proximal tubule epithelial cells (PTECs); x15,000 magnification; scale bar = 1 µm; n = 3-4 rats/group and 5-15 images/rat. Bold solid lines represent the median; lighter dashed lines represent the 25 and 75 percentile (interquartile range). Outliers were first removed using the ROUT test with Q = 1%; normality was tested with a Shapiro-Wilk test; data were analyzed by one way ANOVA followed by Tukey post hoc test. Significance set at $P < 0.05$. NG: normoglycemia; MHG: moderate hyperglycemia; SHG: severe hyperglycemia.

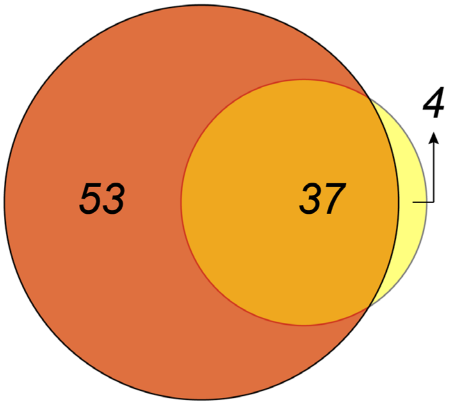


| SHG *vs.* NG only proteins | | | | SHG *vs.* NG & SHG *vs.* MHG overlapping proteins | | SHG *vs.* MHG only proteins | |
|---|---|---|---|---|---|---|---|
| HMGCS2 | COQ7 | ABCB6 | DNM1L | ADHFE1 | COX6B1 | DHODH | HADHA |
| CLPP | SLC25A20 | HADH | SLC25A40 | SLC25A21 | LETM1 | PPOX | CRAT |
| ECH1 | SIRT5 | ECSIT | SLC25A16 | PRODH | SYNJ2BP | | |
| ECI1 | HMGCL | LACTB | DUT | ACSS1 | SLC25A13 | | |
| DECR1 | | NDUFS3 | OCIAD2 | AASS | MFN2 | | |
| MAOB | | SLC25A42 | PPM1K | NADK2 | MICU3 | | |
| ACO2 | | SLC30A9 | ABCG2 | HDHD3 | SLC9A6 | | |
| CBR4 | | AGK | | GSTZ1 | SLC25A23 | | |
| CPT2 | | FLAD1 | | GLYCTK | BDH1 | | |
| ECI2 | | HRSP12 | | AMACR | SLC25A30 | | |
| ACSL1 | | PTRH2 | | SIRT3 | BCKDK | | |
| MLYCD | | SFXN5 | | AMT | SLC25A4 | | |
| ACOT2 | | SLC25A51 | | MTHFS | ACADSB | | |
| DLST | | SLC25A1 | | COQ4 | SLC25A35 | | |
| COQ9 | | DCXR | | SUCLG2 | GATM | | |
| HSPE1 | | ACADM | | NNT | MRPL43 | | |
| CARS2 | | SLC25A45 | | GPX1 | LDHD | | |
| FECH | | COX5A | | HSD17B8 | | | |
| ETFA | | GOLPH3 | | GOT2 | | | |
| ETFB | | HAGH | | PRDX5 | | | |
| PLGRKT | | SLC25A26 | | | | | |

**Figure S2. Differentially expressed proteins between SHG *vs.* MHG and SHG vs. NG.** Venn diagram representation of the overlap (light orange circle, 37 proteins) between the differentially expressed proteins identified in severe hyperglycemia (SHG) *vs.* normoglycemia (NG) (light and dark orange circle, 90 total proteins) and SHG *vs.* moderate hyperglycemia (MHG) comparison (light orange and yellow circle, 41 total proteins). Differentially expressed proteins between group pairs were determined by unpaired t-test with permutation-based false discovery rate correction. Significance was set at $P > 0.05$.

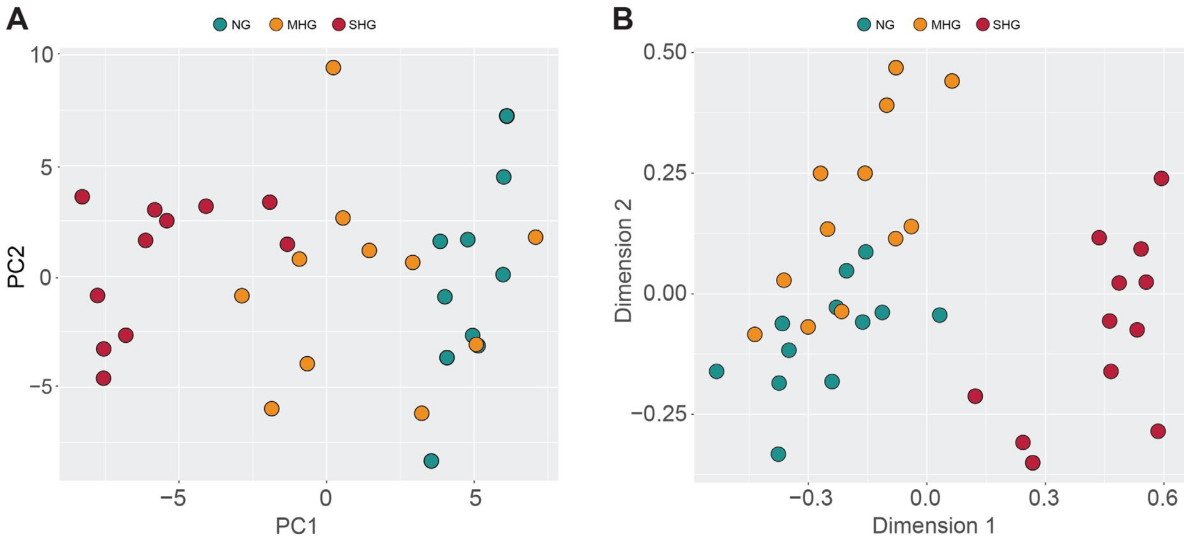


**Figure S3. Principal component analysis (metabolomics) and multidimensional scaling analysis (lipidomics) showing the three treatment groups.** (A) Principal component analysis generated from the metabolomics assessment and (B) multidimensional scaling analysis generated from the lipidomics assessment showing clustering of the severe hyperglycemia (SHG) from the moderate hyperglycemia (MHG) and normoglycemia (NG) groups.

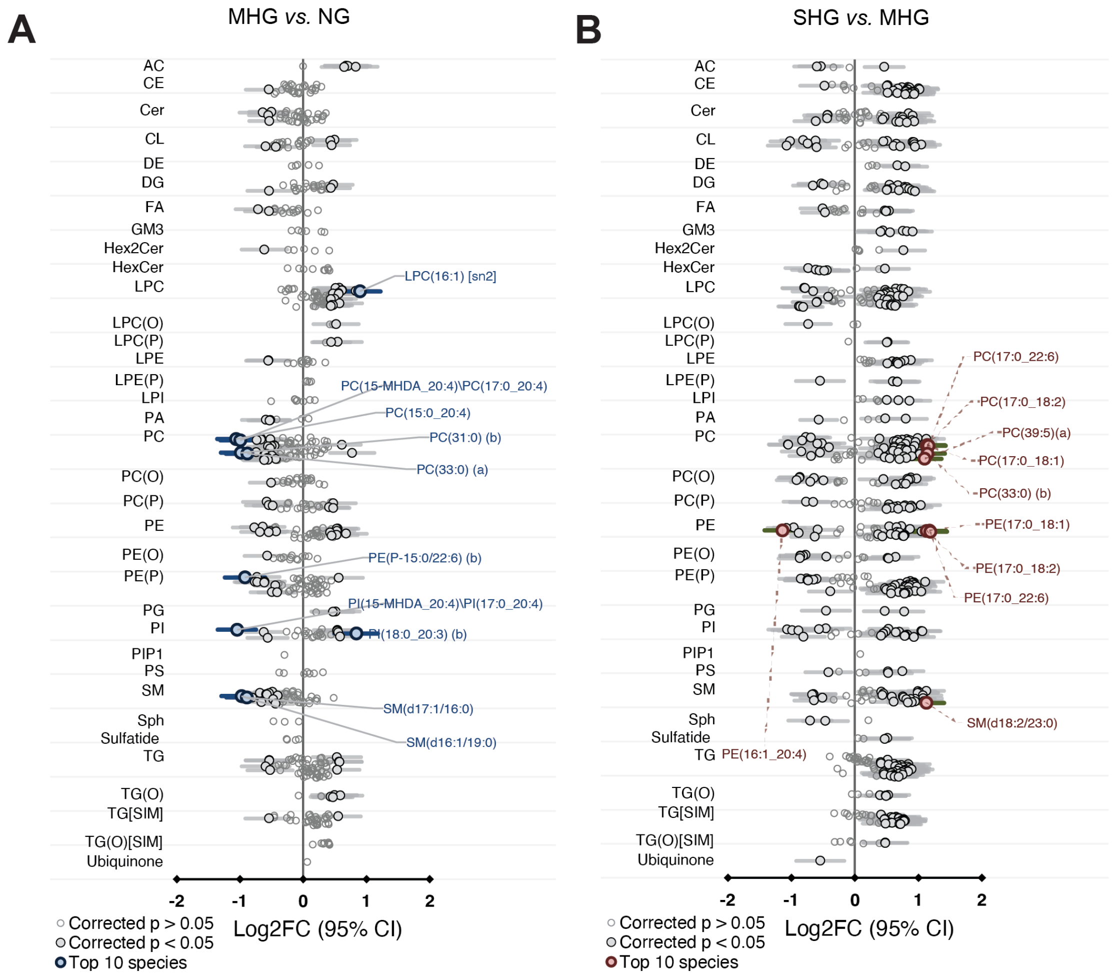


**Figure S4. Lipid species differences between groups.** Forrest plots displaying $Log_2FC$ differences between (A) MHG *vs.* NG and (B) SHG *vs.* MHG in isolated mitochondrial fractions from kidney cortex. Significance set at $P$ = 0.05 after correction for multiple comparisons (Benjamini-Hochberg). Data were normalized by total lipid content. n = 12 for MHG and NG and 11 for SHG. NG: normoglycemia; MHG: moderate hyperglycemia; SHG: severe hyperglycemia.

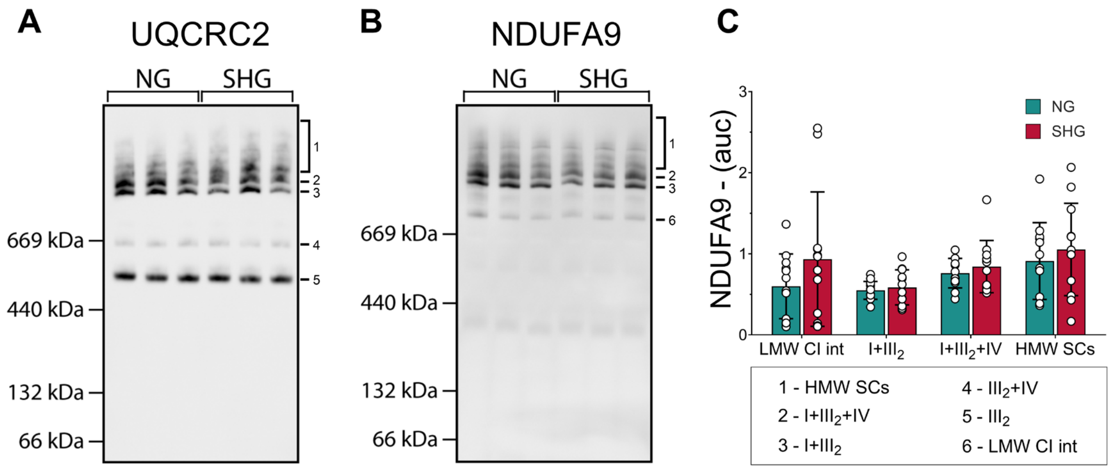


**Figure S5. Cardiolipin remodeling perturbs supercomplex assembly and stability.** Representative BN-PAGE immunoblot image of isolated mitochondria (IM) fractions from rat kidney cortex obtained using an antibody against A) ubiquinol-cytochrome *c* reductase core protein 2 (UQCRC2) and B) NADH:ubiquinone oxidoreductase subunit A9 (NDUFA9); C) quantification analysis of supercomplexes from BN-PAGE immunoblots against NDUFA9. Band 1 (HMW SCs): high molecular weight supercomplexes consisting of complex $I+III_n+IV_n$; band 2 ($I+III_2+IV$): a supercomplex consisting of CI, a dimer of CIII and CIV; band 3 ($I+III_2$): a supercomplex consisting of CI and a dimer of CIII; band 4 ($I+III_2$): a supercomplex consisting of a dimer of CIII and CIV; band 5 ($CIII_2$): CIII dimer; band 6 (LMW CI int): low molecular weight intermediate of CI. n = 12/group. Outliers were first removed using the ROUT test with Q = 1%; normality was tested with a Shapiro-Wilk test; data were analyzed by one-way ANOVA followed by Tukey post hoc test. Data are mean ± SEM. Dots represent individual data points. Significance set at $P < 0.05$. NG: normoglycemia; SHG: severe hyperglycemia.

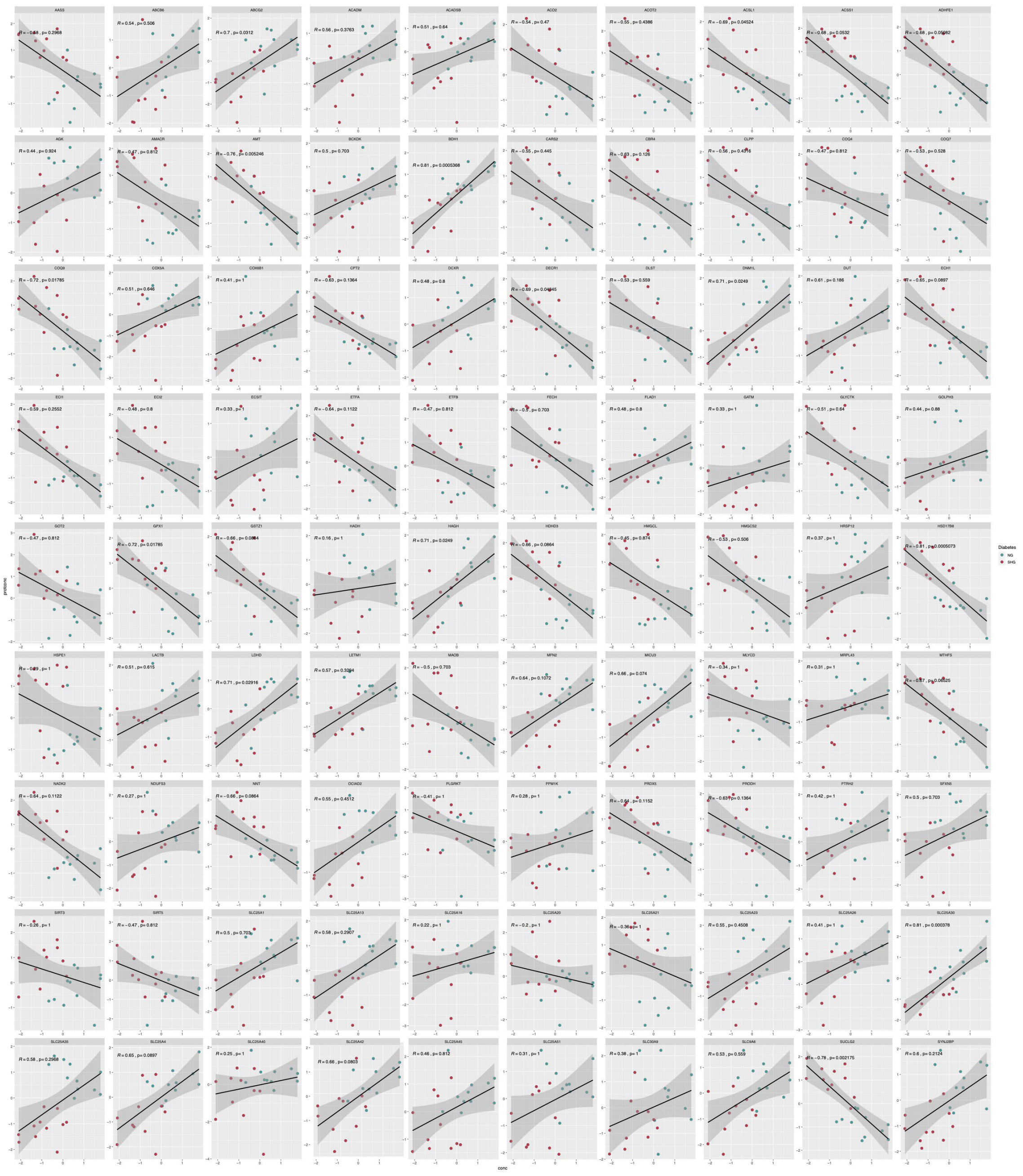

**Figure S6. Correlation plots between plasma cystatin C and all differentially expressed proteins in SHG *vs.* NG.** Correlation plots displaying all correlations (significant and non-significant) between plasma cystatin C and all the differentially expressed proteins between severe hyperglycemia (SHG) and normoglycemia (NG). Plasma cystatin C values (x-axis) underwent a log2 transformation followed by scaling and were correlated against normalized log2 transformed intensity values for proteins (y-axis). Correlation coefficients were calculated using the Spearman method with Holm corrections to account for multiple comparisons. Correlations were considered significant at $P < 0.01$. Dots represent individual values.

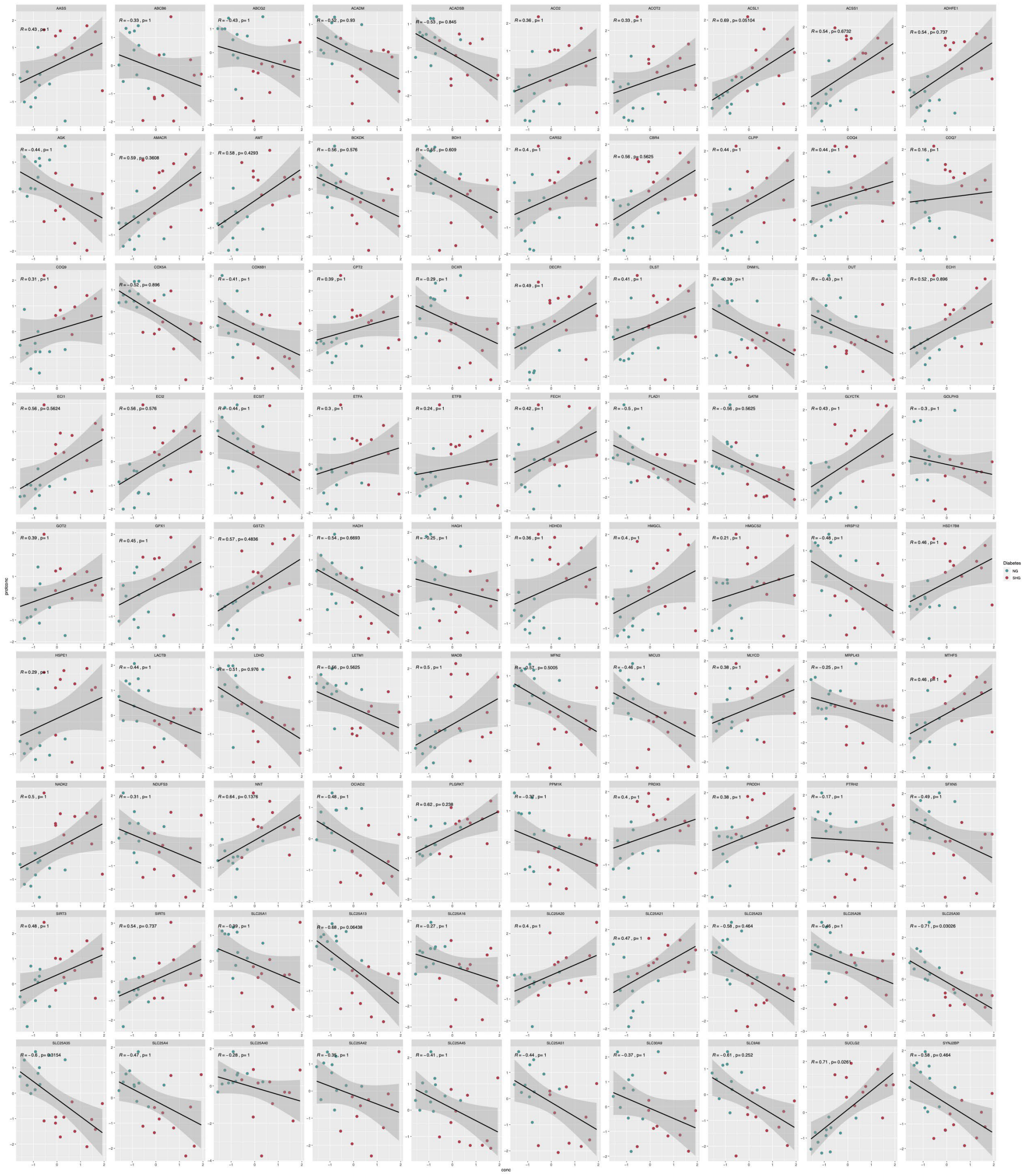

**Figure S7. Correlation plots between urinary albumin and all differentially expressed proteins in SHG *vs.* NG.** Correlation plots displaying all correlations (significant and non-significant) between urinary albumin and all the differentially expressed proteins between severe hyperglycemia (SHG) and normoglycemia (NG). Urinary albumin values (x-axis) underwent a log2 transformation followed by scaling and were correlated against normalized log2 transformed intensity values for proteins (y-axis). Correlation coefficients were calculated using the Spearman method with Holm corrections to account for multiple comparisons. Correlations were considered significant at $P < 0.01$. Dots represent individual values.

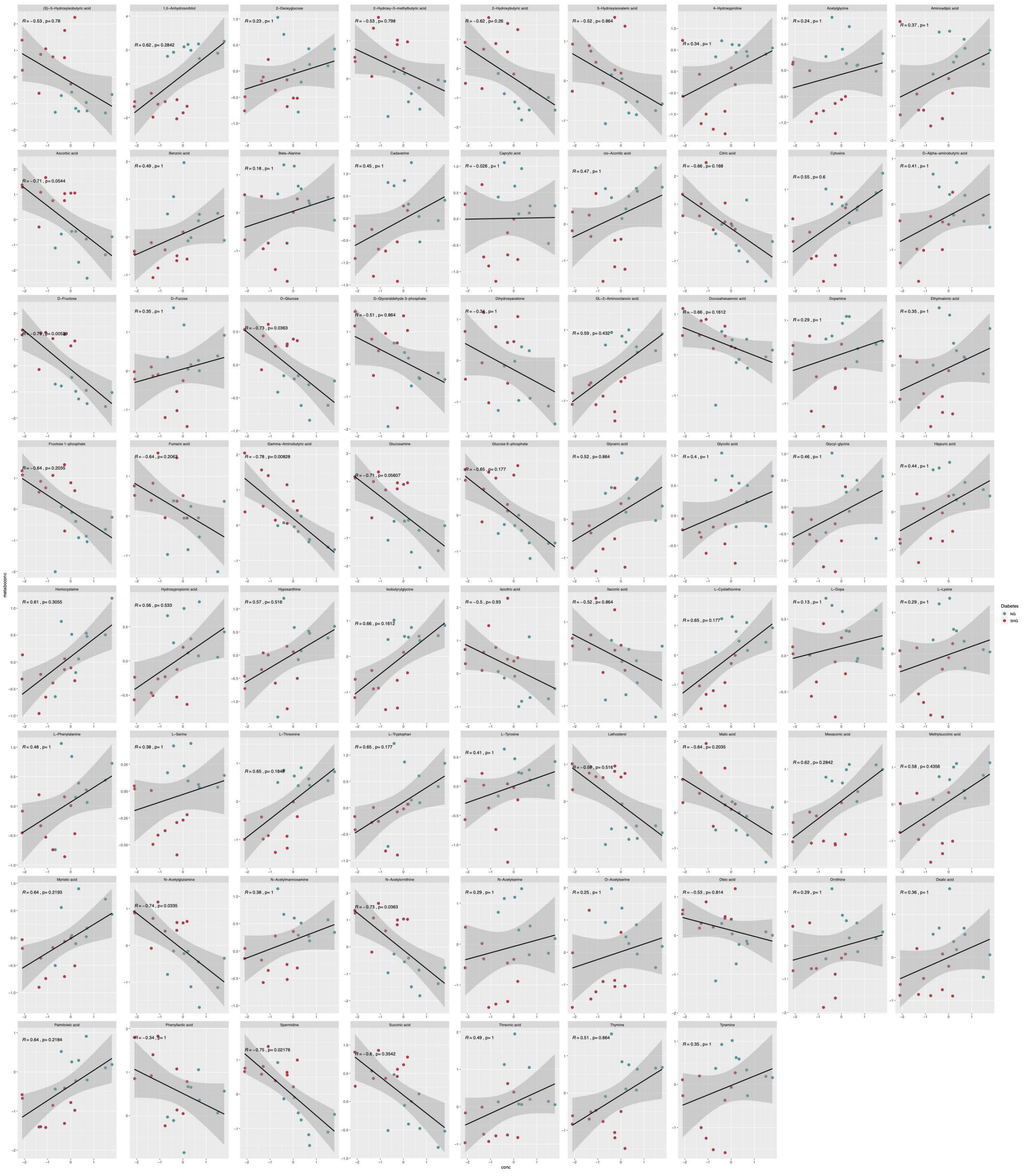

**Figure S8. Correlation plots between plasma cystatin C and all differentially expressed metabolites in SHG *vs.* NG.** Correlation plots displaying all correlations (significant and non-significant) between plasma cystatin C and all the differentially expressed metabolites between severe hyperglycemia (SHG) and normoglycemia (NG). Plasma cystatin C values (x-axis) underwent a log2 transformation followed by scaling and were correlated against normalized log2 transformed intensity values for metabolites (y-axis). Correlation coefficients were calculated using the Spearman method with Holm corrections to account for multiple comparisons. Correlations were considered significant at $P < 0.01$. Dots represent individual values.

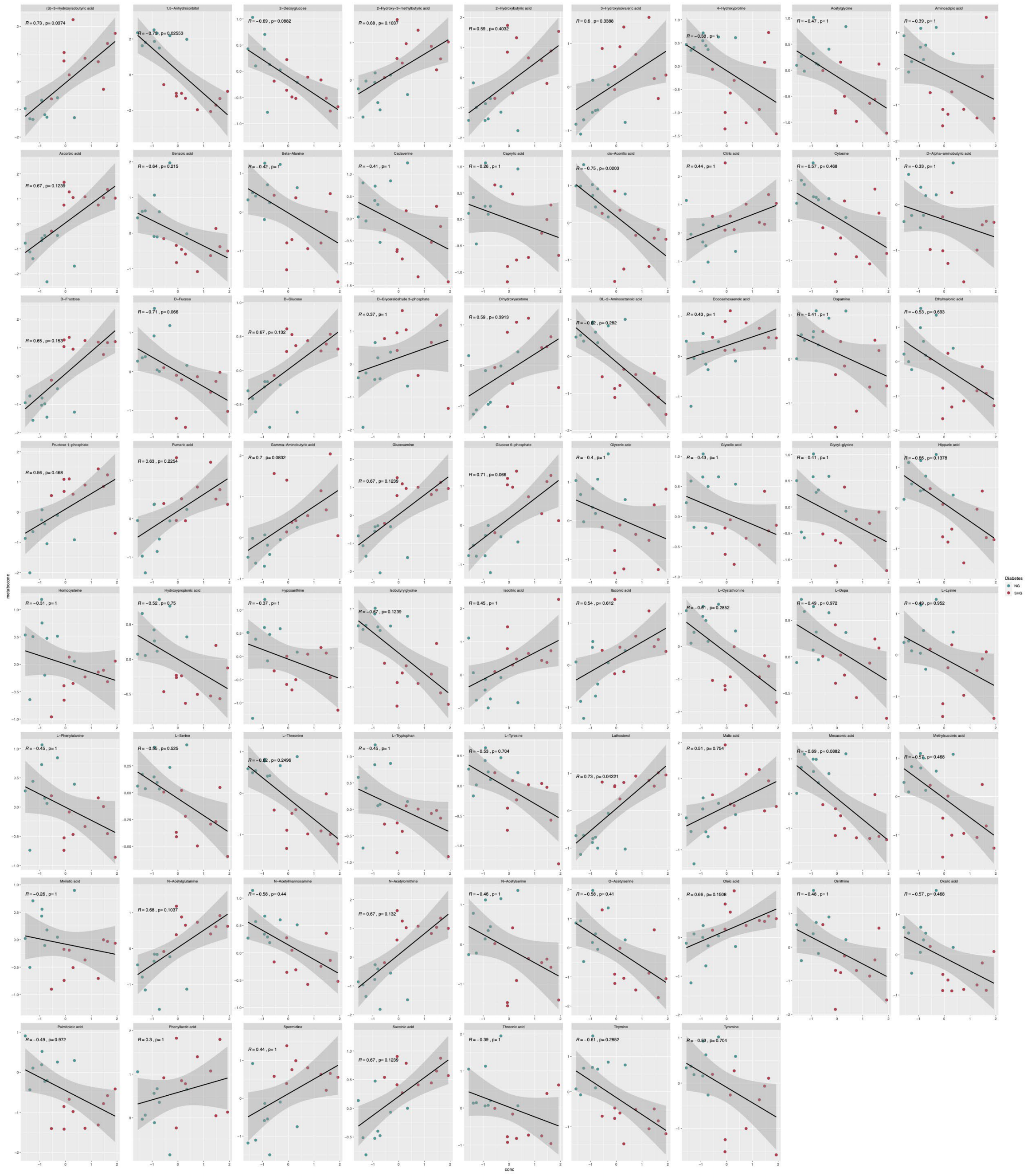

**Figure S9. Correlation plots between urinary albumin and all differentially expressed metabolites in SHG *vs.* NG.** Correlation plots displaying all correlations (significant and non-significant) between urinary albumin and all the differentially expressed metabolites between severe hyperglycemia (SHG) and normoglycemia (NG). Urinary albumin values (x-axis) underwent a log2 transformation followed by scaling and were correlated against normalized log2 transformed intensity values for metabolites (y-axis). Correlation coefficients were calculated using the Spearman method with Holm corrections to account for multiple comparisons. Correlations were considered significant at $P < 0.01$. Dots represent individual values.

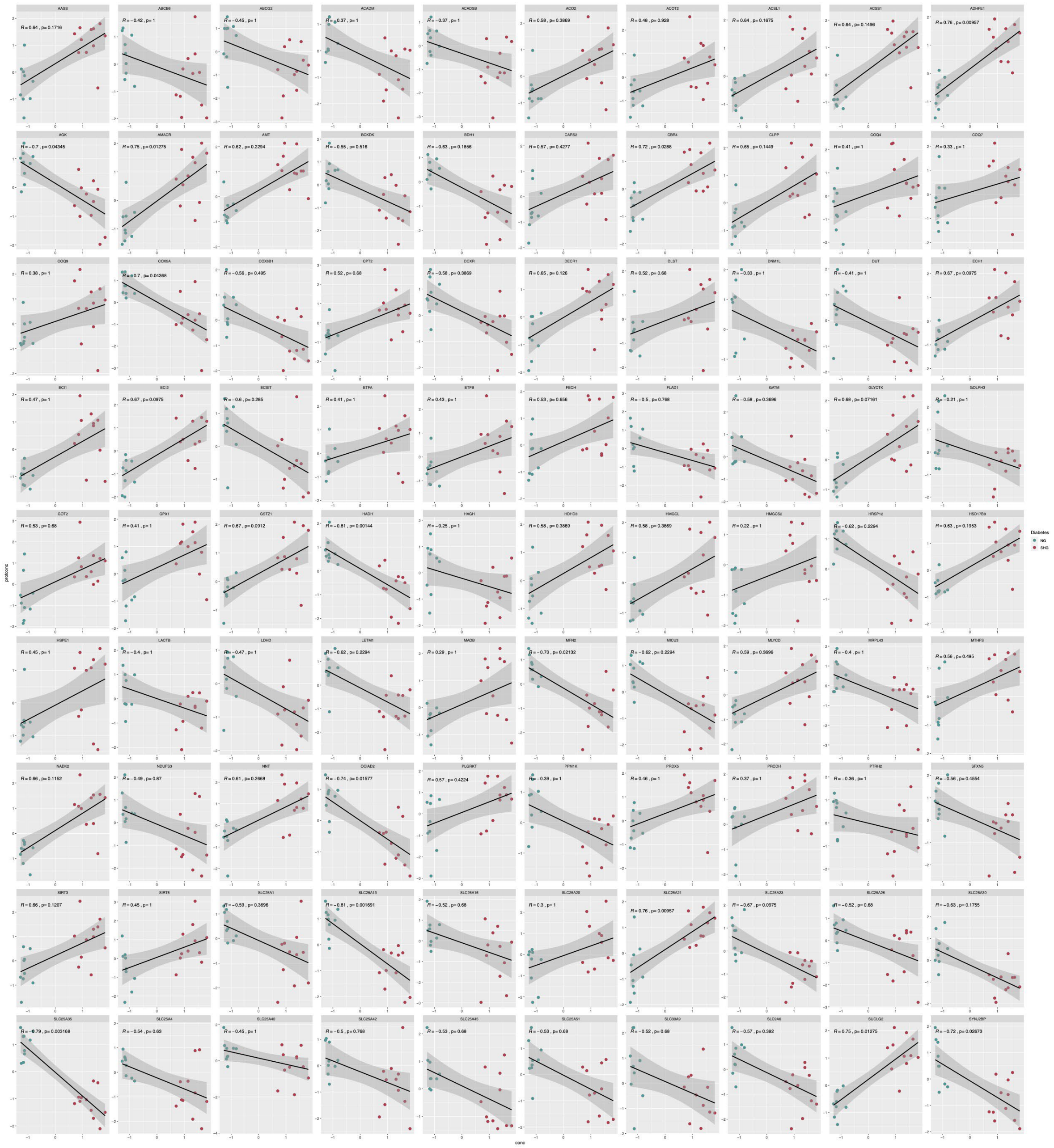

**Figure S10. Correlation plots between HbA1c and all differentially expressed proteins in SHG *vs.* NG.** Correlation plots displaying all correlations (significant and non-significant) between HbA1C and all the differentially expressed proteins between severe hyperglycemia (SHG) and normoglycemia (NG). HbA1c values (x-axis) underwent a log2 transformation followed by scaling and were correlated against normalized log2 transformed intensity values for proteins (y-axis). Correlation coefficients were calculated using the Spearman method with Holm corrections to account for multiple comparisons. Correlations were considered significant at $P < 0.01$. Dots represent individual values.

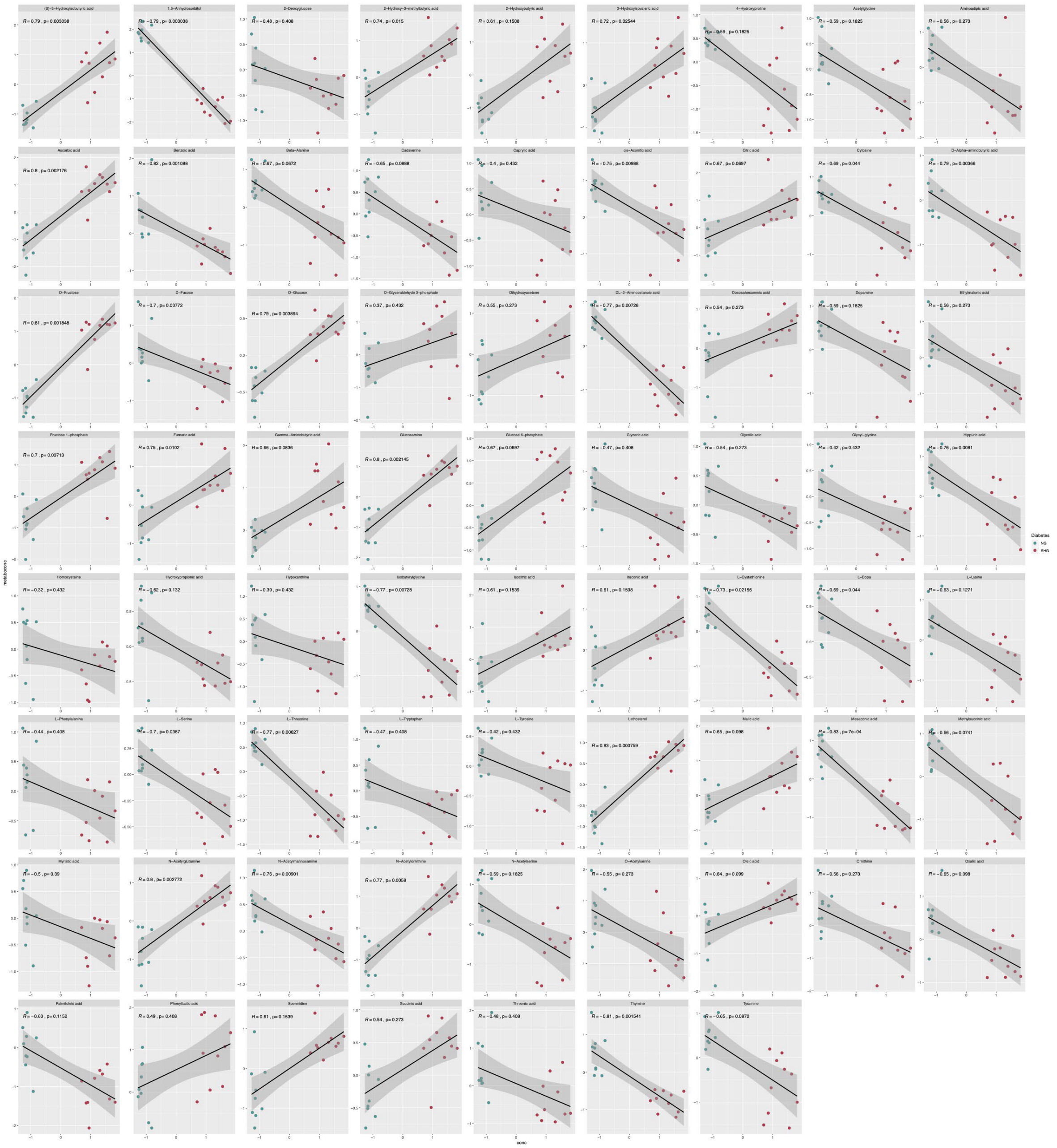

**Figure S11. Correlation plots between HbA1c and all differentially expressed metabolites in SHG *vs.* NG.** Correlation plots displaying all correlations (significant and non-significant) between HbA1C and all the differentially expressed metabolites between severe hyperglycemia (SHG) and normoglycemia (NG). HbA1c values (x-axis) underwent a log2 transformation followed by scaling and were correlated against normalized log2 transformed intensity values for metabolites (y-axis). Correlation coefficients were calculated using the Spearman method with Holm corrections to account for multiple comparisons. Correlations were considered significant at $P < 0.01$. Dots represent individual values.
